# Impacts of the Toba eruption and montane forest expansion on diversification in Sumatran parachuting frogs (*Rhacophorus*)

**DOI:** 10.1101/843664

**Authors:** Kyle A. O’Connell, Jamie R. Oaks, Amir Hamidy, Kyle J. Shaney, Nia Kurniawan, Eric N. Smith, Matthew K. Fujita

## Abstract

Catastrophic events, such as volcanic eruptions, can have profound impacts on the demographic histories of resident taxa. Due to its presumed effect on biodiversity, the Pleistocene eruption of super-volcano Toba has received abundant attention. We test the effects of the Toba eruption on the diversification, genetic diversity, and demography of three co-distributed species of parachuting frogs (Genus *Rhacophorus*) on Sumatra. We generate target-capture data (∼950 loci and ∼440,000 bp) for three species of parachuting frogs and use these data paired with previously generated double digest restriction-site associated DNA (ddRADseq) data to estimate population structure and genetic diversity, to test for population size changes using demographic modelling, and to estimate the temporal clustering of size change events using a full-likelihood Bayesian method. We find that populations around Toba exhibit reduced genetic diversity compared with southern populations, and that northern populations exhibit a shift in effective population size around the time of the eruption (∼80 kya). However, we infer a stronger signal of expansion in southern populations around ∼400 kya, and at least two of the northern populations may have also expanded at this time. Taken together, these findings suggest that the Toba eruption precipitated population declines in northern populations, but that the demographic history of these three species was also strongly impacted by mid-Pleistocene forest expansion during glacial periods. We propose local rather than regional effects of the Toba eruption, and emphasize the dynamic nature of diversification on the Sunda Shelf.

## INTRODUCTION

Catastrophic events can shape patterns of species diversification, population genetic structure, and local demography (Schoener & Spiller, 2006; Botting, 2016). Volcanic eruptions in particular can have profound effects on species’ evolutionary histories, and these events often leave signals on the genomic patterns of resident species (Gübitz et al., 2005; Crisafulli et al., 2015). One Quaternary volcanic eruption has received exceptional attention and controversy due to its presumed impact on biodiversity: the Toba ‘super-eruption’ (Chester et al., 1991; Rampino & Self, 1992; Yost et al., 2018). The Toba eruption occurred ∼74 kya and was the world’s largest Quaternary explosive eruption (Ninkovich et al., 1978; Oppenheimer, 2002; Williams, 2011). Although the Toba Caldera experienced four major eruptions beginning at 1.2 Ma (Chesner et al., 1991), the final eruption was 3,500 times greater than the largest eruption of the modern era (1815 eruption of Tambora) and ash deposits have been identified as far away as India and the South China Sea (Fig. 1A; Rose & Chesner, 1987; Acharyya & Basu, 1993; Westgate et al., 1998; Bühring & Sarnthein, 2000; Oppenheimer, 2002; Robock et al., 2009). Nonetheless, the eruption’s impact on biological life has been extensively debated, with some sources hypothesizing the onset of a global winter that decreased Earth’s temperature by as much as 3– 5°C (Rampino & Self, 1992), leading to dry conditions and pronounced deforestation (Robock et al., 2009; Williams et al., 2009; but see Haslam & Petraglia, 2010). This climate event may have driven local bottlenecks and even extinction (Robock et al., 2009), and has been implicated in precipitating human diversification (Ambrose 1998; 2003; Rampino & Self, 1992).

**Figure 1:**
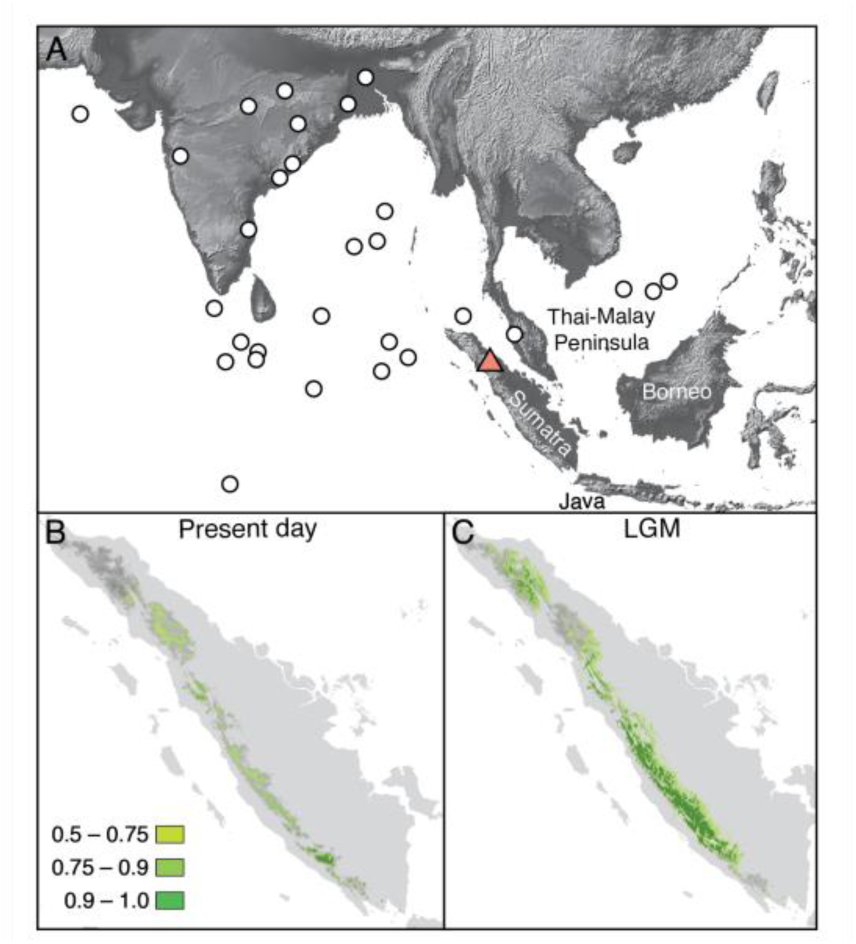
Maps of study region outlining alternative diversification hypotheses. A) Map of south and southeast Asia showing the distribution of recovered ash-fall deposits (grey circles) from the final Toba eruption ∼74 kya (adapted from Oppenheimer, 2002), which was the world’s largest Quaternary explosive eruption. The eruption site of Toba is shown by the red triangle in northern Sumatra. The four landmasses of the Sunda Shelf are also labelled. B) Hypothesized montane forest cover for present day based on species distribution modelling (SDM) of montane forest restricted parachuting frogs (Genus *Rhacophorus*). Present day distributions approximate distributions during Pleistocene interglacial periods. C) Hypothesized montane forest cover at the last glacial maxima (LGM), suggesting that montane forest habitat occupied a greater proportion of the Sumatran highlands during Pleistocene glacial periods than during interglacials.

Alternatively, studies of non-human animals, as well as some human studies, support mixed effects of the eruption on species diversification (Erwin & Vogel, 1992; Gathorne-Hardy & Harcourt-Smith, 2003; Yost et al., 2018). The eruption has been implicated in a wide variety of evolutionary scenarios, including decreased gene flow in orangutans (Nater et al., 2011; 2017), diversification through habitat fragmentation in tigers (Luo et al., 2004), facilitating secondary contact in Sumatran elephants (Fleischer et al., 2001), and driving one haplotype group in palm civets to extinction (Patou et al., 2010). Alternatively, the fossil record provides evidence for few if any extinction events in mammals (Louys, 2007; Louys & Meijaard, 2010. In other volcanic systems, signals of population size change following eruptions have been detected in genetic data supporting both population expansion (Gübitz et al., 2005; Brown et al., 2017) and contraction (Carson et al., 1990; Beheregaray et al., 2003). However, the impact of the Toba eruption on population demography has received less attention than its effect on extinction.

In addition to active volcanism, the island of Sumatra experienced a complex geological history during the Quaternary driven by glacial cycles, which led to periodic connectivity with the rest of Sundaland (Java, Borneo, Thai-Malay Peninsula) and shifts in forest distribution (Fig. 1B,C; Cannon et al., 2009; Hall, 2009; 2012a; 2012b; Lohman et al., 2011; Iwanaga et al., 2012; Morley, 2012; Raes et al., 2014). These climate fluctuations repeated approximately every 100 kya during the Pleistocene (Voris, 2000; Woodruff, 2010; Husson et al., 2019; Sarr et al., 2019). Although lowland forest habitat likely expanded during warmer and wetter interglacial periods, (Morley & Flenley 1987; Heaney, 1991; Bird et al., 2005), cooler climates during glacial periods (2–6 °C lower than today; Van der Kaars & Dam, 1995; Morley, 2012) likely facilitated the expansion of montane forests (Newsome & Flenley, 1988; Stuijts et al., 1984, 1988; Cannon 2009; 2012). Thus, montane forest restricted species may have experienced dynamic population sizes during glacial periods and this signal should be detectable in the demographic history of resident species (Mays et al., 2018).

Despite considerable effort to understand the effects of the Toba eruption and glacial cycles on many species in the region, few studies have investigated how these effects impacted the demographic processes of montane-restricted taxa. One taxonomic group that can be used to address these questions are the parachuting frogs (genus *Rhacophorus*) because they are species-rich and widely distributed across Sumatra. Fifteen species of *Rhacophorus* have been described from Sumatra, and a high proportion are endemic species that diversified *in situ* on the island (Harvey et al., 2002, Streicher et al., 2012, 2014, Hamidy & Kurniati, 2015, O’Connell et al., 2018a,b). In particular three species, *R. catamitus*, *R. modestus*, and *R. poecilonotus*, are co-distributed across the island and are restricted to montane forest streams, allowing for the comparison of populations with similar life histories located proximate to, and distant from the Toba eruption. Past work with these species found evidence of congruent population structure and synchronous divergence (∼5 Ma) across Sumatra, suggesting similar responses to shared historical processes (O’Connell et al., 2018a). However, the same study showed that *R. catamitus* and *R. modestus* originated in northern Sumatra, while *R. poecilonotus* originated in southern Sumatra, suggesting differential responses to some geological events between *R. poecilonotus* and the other two species.

In this study we use target-capture and double digest restriction-site associated DNA (ddRADseq) data to investigate the demographic and evolutionary effects of the Toba eruption and shifts in montane forest distribution on three *Rhacophorus* species from Sumatra. We test the following hypotheses: (1) If the Toba eruption impacted the demographic history of parachuting frogs, we expect that (a) northern populations will exhibit reduced genetic diversity compared with southern populations (b) analyses will support population contraction in the northern populations around the time of the eruption, as documented in other volcanic systems. (2) If cycles of forest expansion and contraction impacted the demographic history of parachuting frogs, we expect that (c) genetic diversity will show no geographic pattern across the island, (d) northern and southern populations will exhibit congruent timing of size change events corresponding to discrete glacial cycles.

## MATERIALS AND METHODS

### Species distribution modelling

We used species distribution models to infer the extent of montane forest during glacial and interglacial periods. We compiled locality information for the three focal species: *R. catamitus*, *R. modestus*, and *R. poecilonotus* during our field surveys conducted across Sumatra in 1996 (Harvey et al., 2002) and from 2012-2016 (Harvey et al., 2015, 2016). In an effort to understand changes in species distributions during glacial cycles, we estimated species distribution models using MaxEnt 3.3.3k (Phillips et al., 2006). Specifically, we used 19 bioclimatic variable files at 30s resolution for contemporary climate data, and 19 variables at 2.5m resolution for Last Glacial Maximum (LGM) data (highest resolution available) from WorldClim (2020). Although the LGM likely represents a more severe climate fluctuation than most of the Pleistocene glacial cycles experienced on Sundaland (Morley, 2012), it is the only glacial cycle with suitable data and thus serves as a useful approximation of montane forest extent during glacial cycles. Likewise, we use contemporary distributions of the three focal species to understand species distributions during interglacial periods. We noted no microhabitat differences between the focal species during field work, and thus, grouped all species into a single model to increase sample sizes and model the distribution of the species assemblage through time. We formatted bioclim files using the *Clip* and *Raster* to ASC tools in QGIS 3.6 (available from available at https://github.com/qgis/QGIS). In MaxEnt, we applied the auto features settings, and applied 5,000 iterations. We changed the replicated run type to ‘Subsample’ and assigned the random test percentage to 25 and used the *Mask* tool in QGIS to crop out zones that fall outside our study area. We evaluated model performance by assessing the omission data plot and area under the curve (AUC) plot. The resulting SDM models indicated the probability of occurrence (0.0 – 1.0) for the assemblage, which should serve as a good proxy for montane forest extent given the habitat specificity of our target species. All layers used in our analyses are available online (Data Accessibility).

### Genomic data generation

Genomic data were generated followed Portik et al., (2016) with modifications. Exonic targets were designed from transcriptome sequences from *R. modestus* and *R. monticola* using custom scripts (see Data Accessibility). Probes for the target sequences were synthesized by myBaits® (Arbor Biosciences, Ann Arbor, MI) using 120 bp probes with 2x tiling. The final probe set resulted in 19,950 baits targeting 955 transcript markers from each of the two species. Library preparation was conducting using the Kapa Hyper Prep Kit (Kapa Biosystems, Wilmington, MA) for 69 individuals. Starting with 1 μg of DNA from each pooled library group, we hybridized pooled genomic libraries with target probes following the myBaits® protocol. We pooled capture libraries into two sequencing groups (Table S1) which were sequenced on two lanes (150 bp PE) of the Illumina® X10 at Medgenome (medgenome.com). Full methods for transcriptome data generation and assembly, probe design, library preparation, and hybridization reactions are described in the Supporting Information.

### Genomic data processing

Phased alignments for downstream analyses were generated using the Python pipeline SECAPR v.1.14 (Andermann et al., 2018) following the steps outlined in the developer documentation (https://github.com/AntonelliLab/seqcap_processor) and fully described in the Supporting Information. We further filtered the final phased alignment using custom Python scripts. We refer to this dataset as ‘full-locus’ because it includes the target exons as well as assembled flanking sequence. All scripts used in filtering are available online (see Data Accessibility).

In order to optimize our alignments at the phylogenetic level (across the genus) and also to evaluate the utility of only exonic sequence for population genomic analyses, we generated a second dataset that we refer to as the ‘exonic’ dataset. Rather than assembling reads then matching the assembled contigs to a reference sequence, we mapped our cleaned reads directly to our target sequences, then phased, aligned, and filtered these alignments as described above and in the Supporting Information.

We called SNPs for both datasets using the *R* script fasta2vcf.R (https://github.com/gehara). To call SNPs for the exonic dataset we first separated our alignments by species so that SNPs were only retained within each species; the full-locus dataset was already separated by species. We report the total number of SNPs retained, the number of loci with SNPs, and the percent missing data in Table 1. Missing data was calculated following de Medeiros and Farrell (2018).

**Table 1:** Summary statistics of target capture datasets for each species. Abbreviations are as follows: Total bp = length of alignment, Seqs/aln = the number of individual samples included in the alignment, % missing = the percent missing data for each alignment, Loci/SNP = the number of loci with a variant site for that species.

| Species | Dataset | Total bp | # Loci | Seqs/aln | Mean Length (bp) | % missing | Total SNPs | Loci w/ SNP | SNPs/locus |
| --- | --- | --- | --- | --- | --- | --- | --- | --- | --- |
| *catamitus* | Full-locus | 250616 | 740 | 10.75 | 338.7 | 25.5 | 2935 | 600 | 4.89 |
|  | Exonic | 440024 | 944 | 11.25 | 466 | 14.25 | 1836 | 488 | 3.76 |
| *modestus* | Full-locus | 241721 | 691 | 7.9 | 349.8 | 15 | 2261 | 530 | 4.27 |
|  | Exonic | 440024 | 944 | 8.1 | 466 | 12.7 | 2336 | 494 | 4.73 |
| *poecilonotus* | Full-locus | 326419 | 711 | 5.95 | 459 | 27 | 2237 | 532 | 4.20 |
|  | Exonic | 440024 | 943 | 6.95 | 466.5 | 15.3 | 1533 | 466 | 3.29 |

**Table 2:**
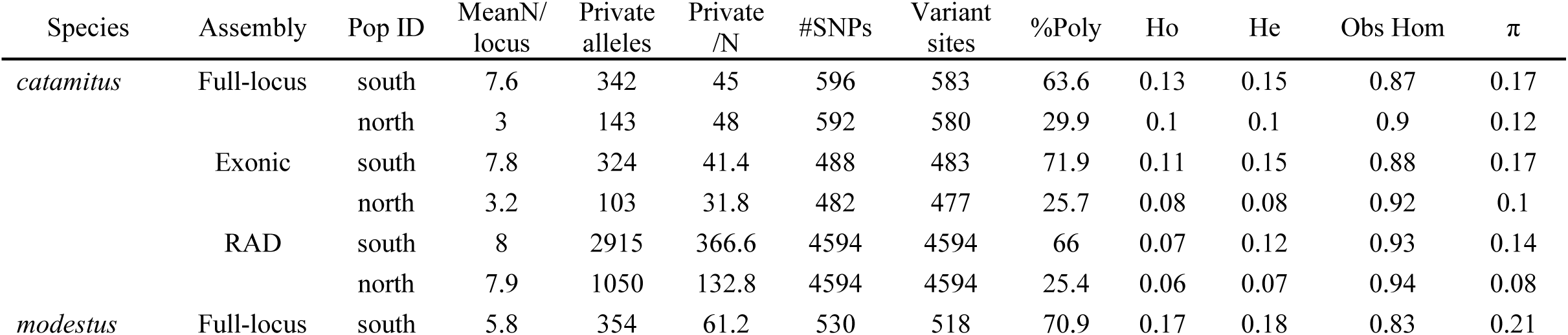
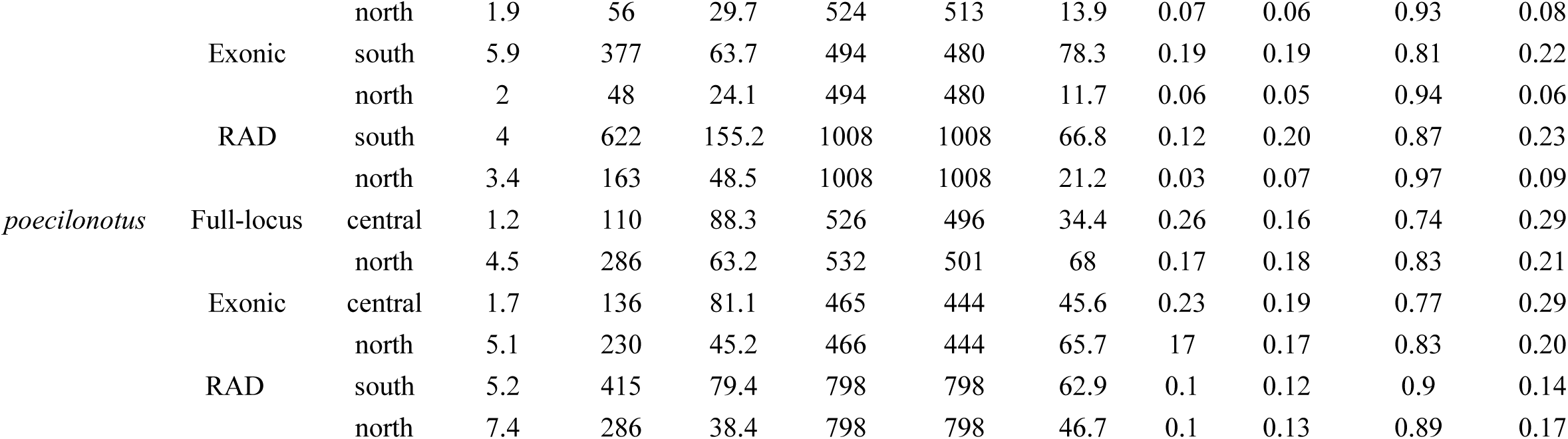
Population-level summary statistics for each species and dataset. Abbreviations are as follows: MeanN/locus = average number of samples in the alignment for a given locus, Private/N = The number of private alleles divided by the average number of individuals per alignment to correct for sampling bias in private allele estimation, %Poly = the percent of sites in a locus that are polymorphic. Values of the following represent mean values across the 500 runs: Ho = observed heterozygosity, He = expected heterozygosity, Obs Hom = observed homozygosity. π = nucleotide diversity.

To compare population genomic parameters estimated using our two target-capture datasets to those estimated from ddRADseq data, we utilized ddRADseq-derived SNP data for all three species from O’Connell et al., (2018a). These datasets were filtered such that 70% of individuals contained a locus for it to be retained. Dataset characteristics were as follows: *R. catamitus* n = 20 with 4594 SNPs and 20.7% missing data; *R. modestus* n = 10 with 1008 SNPs and 26.31% missing data, and *R. poecilonotus* n = 17 with 798 SNPs and 25.5% missing data.

### Testing probe efficacy to recover population structure and estimate genetic diversity

To verify that our capture data recovered the same population structure as mitochondrial and ddRAD data from O’Connell et al., (2018a), we estimated population structure using maximum likelihood using Admixture v.1.3.0 (Alexander et al., 2009) using a range of *K* values (1–5), with five iterations per *K* value. We removed individuals with less than 25% of SNPs, and plotted the mean *K* value for each individual across iterations.

We estimated population genetic summary statistics for each species using SNP alignments from the all datasets using the populations module from STACKS v.2.4 (Catchen et al., 2013), and compared the results using all SNPs and only one random SNP per locus for the two target capture datasets (Table S3). To account for sample size discrepancies between populations and datasets, we implemented a subsampling without replacement approach using custom scripts (Data Accessibility) where we down sampled the vcf file of each population for each species and each dataset for between two samples, which was the minimum number of individuals in a population (*R. poecilonotus* south) to nine samples, which was the maximum number of individuals in a population (*R. catamitus* south). We then ran the populations module on the subsampled datasets for 500 iterations, giving us 500 estimates from a range of sample sizes for each parameter.

Finally, we tested how well our recovered loci could reconstruct the phylogeny using taxa from across the genus. We concatenated all loci for the exonic dataset for all samples, called a consensus sequence for each individual, and estimated a maximum likelihood phylogeny using RAxML v.8.2.11 (Stamatakis, 2014) with 100 rapid bootstrap iterations, partitioning by locus and applying a single GTRCAT model to each partition.

### Demographic modelling of size change events

To investigate if size changes occurred in Sumatran *Rhacophorus* (indicative of catastrophic geological events), we used the diffusion approximation method implemented in δaδi (Gutenkunst, Hernandez, Williamson, & Bustamante, 2009) to analyse two-dimensional Site Frequency Spectra (2D-SFS). The folded 2D-SFS was generated from VCF files (https://github.com/isaacovercast/easySFS), and to account for missing data we projected down to smaller sample sizes for each dataset (Table 3). In the case of the admixed individual of *R. poecilonotus* (see Results), we tested models where we assigned this individual to the northern and southern population. Because we observed only minor differences in parameter estimates, we only present results for this individual assigned to the northern population. For each dataset we compared divergence with no size change and divergence with size change (to estimate expansion or contraction) using the workflow outlined by Portik et al., (2017; https://github.com/dportik/dadi_pipeline). Following Portik et al., (2017) and Barratt et al., (2018), we performed five iterations of each model. Each run consisted of four rounds of optimizations with multiple replicates and used search parameter estimates from the best scoring replicate (highest log-likelihood) to seed searches in the following round. We used the default settings of dadi_pipeline for each round (grid size = 30,40,50; replicates = 10,20,30,40; maxiter = 3,5,10,15; fold = 3,2,2,1), we optimized parameters using the Nelder-Mead approach (optimize_log_fmin), and used the optimized parameter sets of each replicate to simulate the 2D- SFS. The log-likelihood of each 2D-SFS was estimated for each model using a multinominal approach and we identified the best-supported model using log-likelihood and AIC, confirming that the best supported model was consistent across the five iterations. For the target capture datasets we compared models using all SNPs and a single random SNP for each locus; because using linked sites violates the assumptions of AIC model selection (and because results were congruent between datasets), we only report the parameter estimates for the best-supported model from the single SNP analyses (Table 3).

**Table 3:**
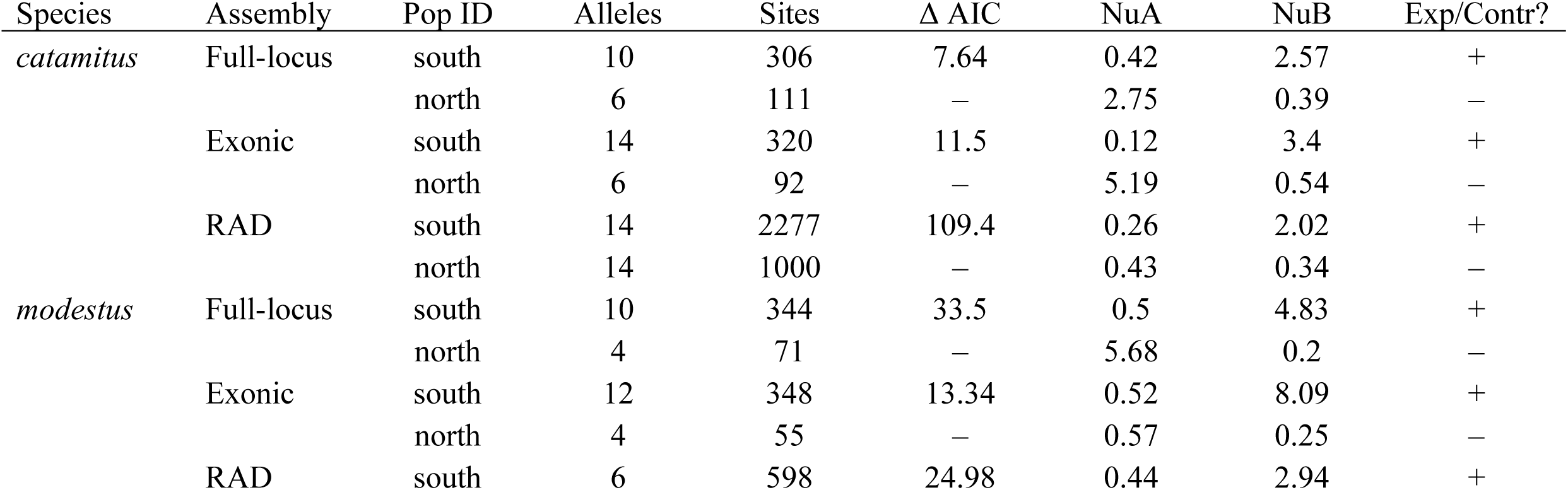
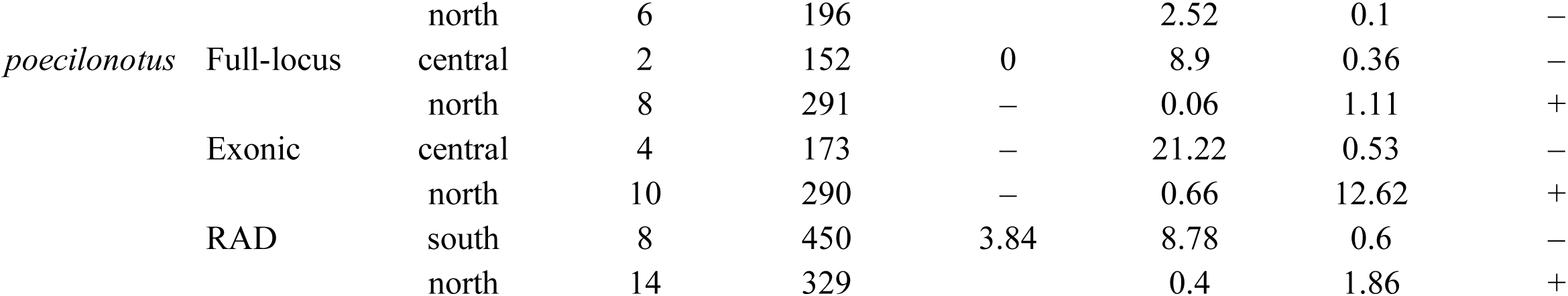
Summary of demographic modeling conducted in δaδi. Alleles and sites refers to the number of alleles and sites included after downscaling, where alleles = twice the number of individuals. Δ AIC refers to the change in AIC value of the best size change model compared to the best model without size change and is only reported once for each population pair as both populations were included in the same model. NuA = scaled population size before size change event, NuB = scaled population size after size change event. + signifies population expansion, while – signifies contraction.

### Estimating the timing of demographic events

We used ecoevolity v.0.3.2 (Bryant et al., 2012; Oaks, 2019a) to estimate the timing and synchronicity of demographic events. Ecoevolity is a general, full-likelihood Bayesian method that can estimate the temporal clustering of demographic events for all populations in a single analysis. The program thus estimates the timing of demographic events, as well as effective population sizes of ancestral and contemporary populations (which are used to infer shifts in population size). To generate input files for each dataset, we included the full alignments for each locus; Oaks et al., (2019b) showed that using all sites substantially improves estimates of demographic event times. We separated each alignment into northern and southern populations for each species based on clustering analyses (Fig. 2) for a total of six populations being compared by ecoevolity. In the case of the admixed *R. poecilonotus* individual we ran analyses once with the individual assigned to each population.

**Figure 2:**
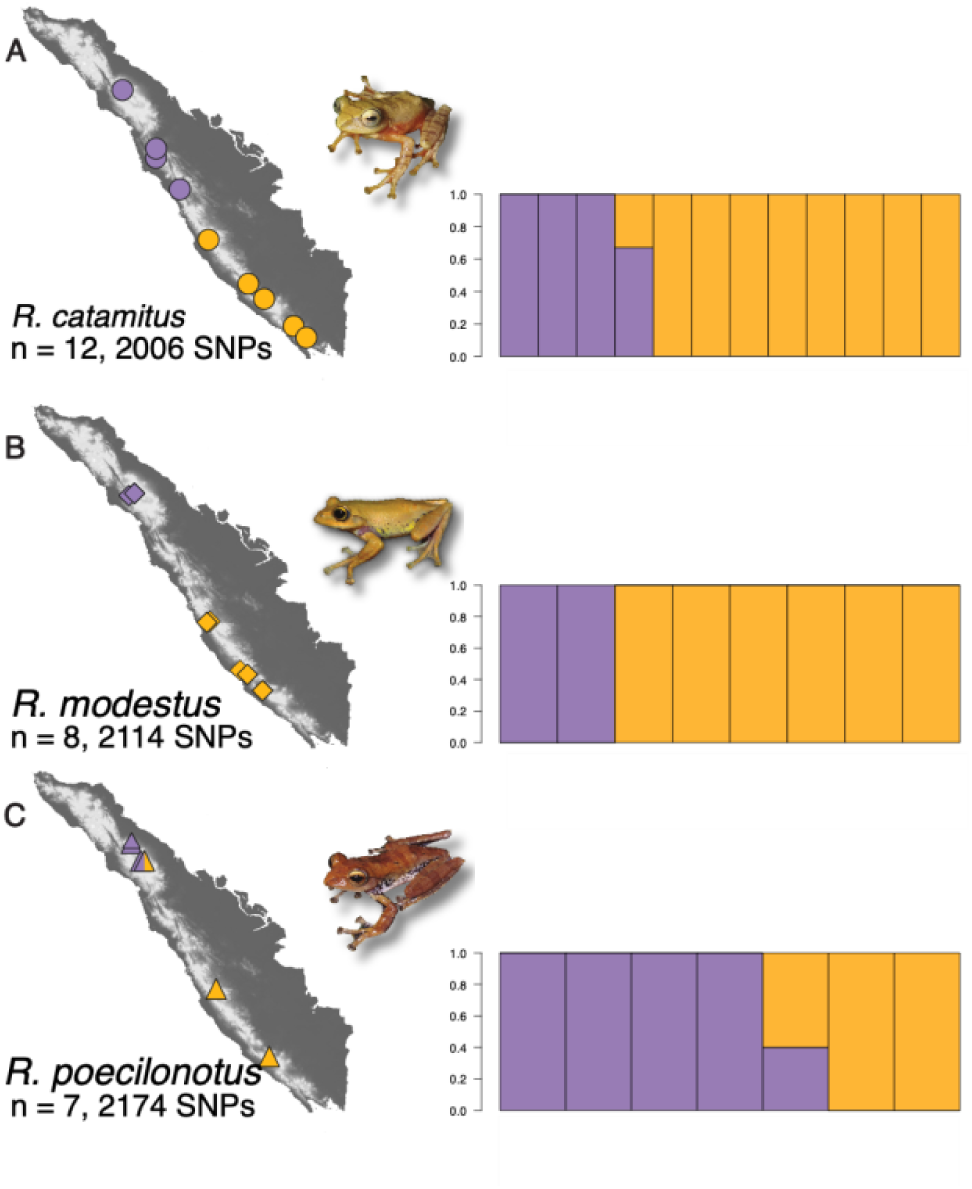
Sampling for the target capture datasets used in this study. The number of SNPs listed reflects the full-locus datasets (see main text). Maps demonstrate the geographic location of each sample, and are accompanied by ancestry proportions estimated in ADMIXTURE v.1.3.0 (Alexander et al., 2009). Individuals from northern populations are coloured purple, southern populations are coloured orange. Maps and ancestry proportion plots correspond to A) *R. catamitus*, B) *R. modestus*, and C) *R. poecilonotus*.

We tested up to 13 parameter combinations (see Supporting Information) and implemented the following parameters in the final analysis: we applied a Dirichlet process with a concentration parameter fixed at 2.62 (mean prior probability on 3.5 demographic events across the six populations) as the prior over all the ways the timing of the population-size change of the six populations can be assigned to 1–6 events, and an exponential distribution for the prior on event times (i.e., unique times of population-size change), with a rate of 1000 (∼500 Kya assuming a mutation rate of 1.42 x 10^-9^ substitutions per site per year). For the prior on the current effective population size (scale by the mutation rate) we used a gamma-distributed prior (shape = 3.5, scale = 0.001). For the prior on the relative effective size of the population before the size change occurred, we used our δaδi results to apply population-specific exponential distributions. On the northern populations of *R. catamitus* and *R. modestus* we applied an exponential distribution with a mean = 2 (modelling contraction), and on the southern populations applied an exponential distribution with a mean = 0.5 (modelling expansion). In *R. poecilonotus* we applied the inverse, and in all cases allowed the parameter to be estimated. We set the mutation rate equal to one and converted posterior estimates by dividing by the mutation rate of 1.42 x 10^-9^ substitutions per site per year, estimated for the frog genus *Leptopelis* (Allio et al., 2017). We ran 20 MCMC chains for each prior combination for each dataset for 75,000 generations with a sample taken ever 50 generations, producing 1500 samples from each chain. We checked for convergence and calculated median values for parameters of interest in Tracer v.1.7.1 (Rambaut et al., 2018) with a 10% burnin. Figures were produced using the accompanying Python package ‘pycoevolity’ v.0.2.4 with a burnin of 101 samples for each log file.

### Simulations to investigate dataset variation

To better understand the variability in posterior estimates between datasets, we used ‘simcoevolity’ v.0.2.4 (Oaks et al., 2019b) within the pycoevolity package to simulate 300 replicates using the ‘-c’ option, which simulates alignments that matched our empirical datasets for numbers of individuals, locus number, locus length, and patterns of missing data within each locus. We also simulated an additional full-locus dataset with double the number of individuals (full-doubled). This strategy allowed us to identify if differences between estimates from the datasets were due to differences in the number of sites, loci sampled, patterns of missing data within each locus, or sample sizes. For the three analyses (exonic, full-locus, and full-doubled), we analysed the 300 simulated replicates using 4 chains of ecoevolity under the parameters described above, and plotted the 4000 MCMC samples for each (1500 samples per chain, less 500 burnin per chain) using ‘pyco-sumsims’ of the pycoevolity package with a burnin of 500 samples per chain.

## RESULTS

### Species distribution modelling

Our species distribution models (SDMs) indicated that during interglacial periods (represented by contemporary distributions), suitable montane forest cover was restricted to upper elevation zones throughout the Barisan Range of western Sumatra (Fig. 1B). Interglacial periods thus likely had regions of non-contiguous montane forest at lower elevations in regions separating montane habitats, suggesting that montane forests are currently in a refugial state (Cannon et al., 2009). In contrast, LGM models supported largely contiguous montane forest across the entire range of the focal species’, spanning many of the lower lying areas (Fig. 1C), providing potential corridors for dispersal of montane taxa.

### Target capture efficiency

We compared two assembly strategies for generating final alignments for the three focal species (Table 1). Generally, the full-locus dataset generated more total SNPs, more SNPs per locus and more loci with SNPs. However, the full-locus dataset had fewer total loci recovered, shorter loci, fewer individuals per alignment, and greater levels of missing data when compared with the exonic dataset. This assembly method was also more sensitive to divergence from the reference sequence, as observed when comparing capture success between *R. modestus* (the reference species) and the other two species (Table 1).

### Estimation of population structure

Within the focal species, the target-capture data recovered similar population structure to the ddRADseq data, with northern and southern populations roughly divided at the midpoint of the island (Figs. 2; S2). In *R. poecilonotus* we inferred one individual with mixed ancestry from northern Sumatra, but this pattern could be caused by either incomplete lineage sorting or secondary contact following allopatric speciation (McGuire et al., 2007; Bell et al., 2015).

### Genetic diversity estimation

Across species, we consistently estimated higher genetic diversity in southern populations compared with northern populations (Fig. 3; S3; Tables 2; S3). Across datasets, parameter values were consistent with the target-capture data when using all SNPs compared with only one SNP (Table S3), and between the full-locus and exonic datasets (Fig. S3; Table 2). In preliminary analyses with unequal sample sizes, we observed large discrepancies between the target capture and ddRAD datasets (results not shown), but when we utilized a bootstrapping approach this discrepancy largely disappeared (Fig. S3). Two exceptions were northern *R. modestus* and southern *R. poecilonotus*, which had smaller sample sizes in the ddRAD datasets.

**Figure 3:**
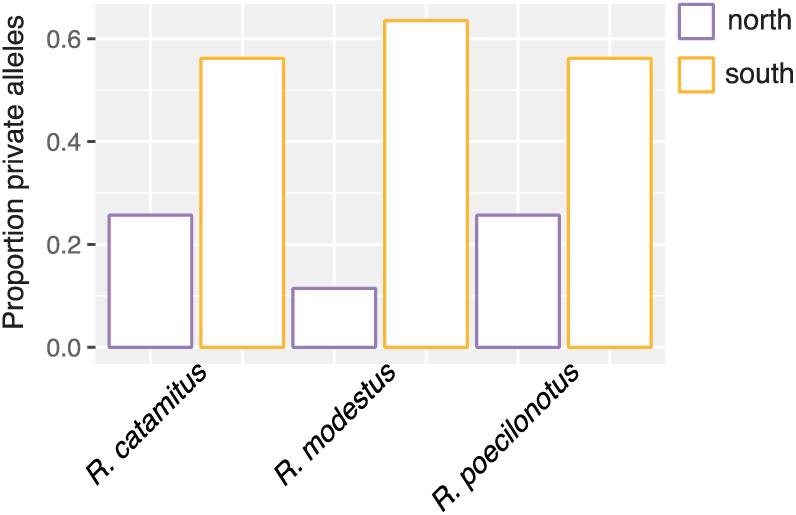
Proportion of private alleles for the full-locus dataset estimated using the Populations module of STACKS v.2.4 (Catchen et al., 2013). Southern populations (coloured orange) exhibit higher levels of genetic diversity than northern populations (coloured purple), consistent with a historical reduction in effective population size in the northern population.

### Demographic model testing

We compared divergence with and without size change for each dataset (Table 3). In all datasets for *R. catamitus* and *R. modestus*, divergence with size change was strongly supported (Δ AIC > 5). However, in *R. poecilonotus*, analyses of the full-locus and exonic datasets either failed to strongly differentiate between models or found weak support for divergence without size change (ΔAIC = 3.8), including in the models where we changed the assignment of the admixed individual (results not shown). In the ddRADseq data for *R. poecilonotus* the size change model was only weakly supported (ΔAIC = 3.8). This suggests that the number of SNPs or the population-level sampling in *R. poecilonotus* (two central individuals and no southern individuals) were not sufficient to differentiate between these two models. We otherwise inferred consistent patterns across analyses (Table 3).

Patterns of population size change were consistent in *R. catamitus* and *R. modestus*, the southern populations always expanded and the northern populations always contracted. In both species the magnitude of southern expansion was greater than that of northern contraction. In *R. poecilonotus* we inferred the opposite signal across all datasets, with northern expansion and southern contraction, and the magnitude of southern contraction was much greater than that of northern expansion. These findings were consistent when including all SNPs or only one SNP, as well as in the ddRADseq data (Table 3).

### Temporal clustering of demographic events

We tested a wide range of priors for each dataset to ensure that our data, and not the prior distributions, were driving posterior estimates (Table 4). We found that the posterior distribution of demographic event times varied little between runs with different priors, but that the prior on the concentration parameter affected support for the number of events between runs, although the best-supported number of events did not vary widely (e.g. 3 vs. 4 events; Figs. S3,4). Nonetheless, we inferred strikingly different results when applying the same priors to different datasets (full-locus vs. exonic; Fig. S4; Table 4). After summarizing results across chains, the ddRAD data failed to converge across the 13 parameter combinations; as such we do not report these posterior estimates. Due to the uncertainty around some of our posterior estimates, as well as the dependence of posterior estimate conversion on an accurate mutation rate, we advise caution when interpreting our posterior estimates in years.

**Table 4:**
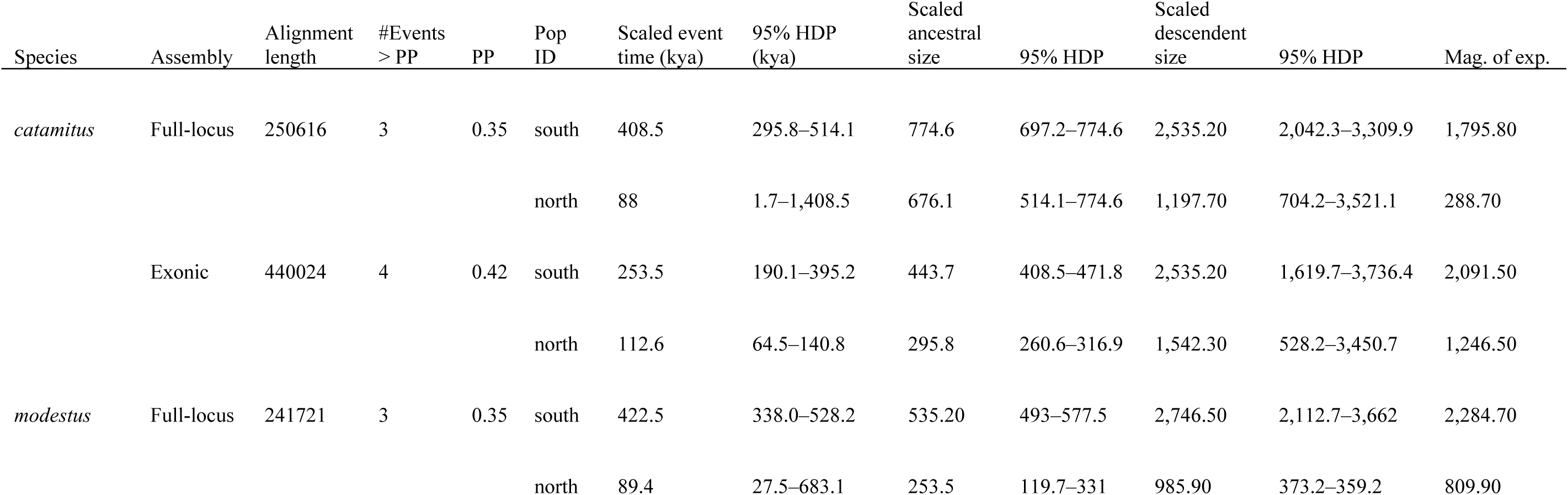
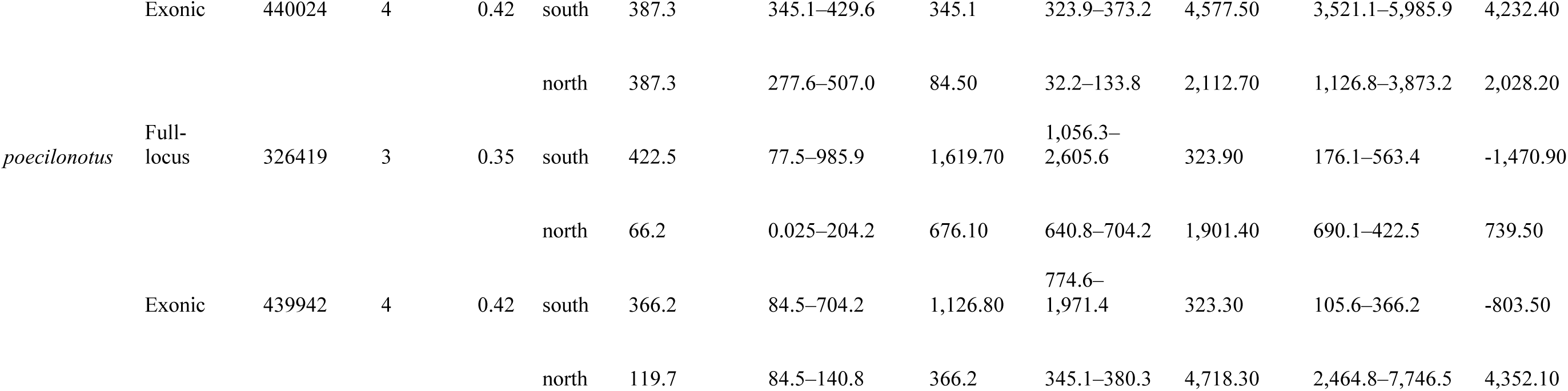
Summary statistics from ecoevolity (Oaks 2019a) analysis estimating temporal clustering of demographic events. Parameters are scaled by the mutation rate of 1.42 x 10^-9^ substitutions per site per year, estimated for the frog genus *Leptopelis* (Allio et al., 2017), and assume a one year generation time. Population sizes, including ancestral, descendent, and magnitude of expansion, are shown in the thousands of individuals. Negative expansion values represent contraction events. Scaled event times show the median values of the posterior distribution, except in the case of northern *R. catamitus* and *R. modestus* which exhibited bimodal distributions; we report the median value of the first peak for both species. The second peak, shown in Figs. 4,S4 is synchronous with the southern populations. Mag. of exp. = magnitude of expansion in thousands of individauls. Negative values represent contraction events.

In the full-locus dataset, three demographic events received the highest posterior probability (PP = 0.35; Fig. S4A). The analysis inferred two general clusters of events, which were consistent with one event shared by the northern populations (∼80 kya), and one event shared by the southern populations (∼400 kya; Fig. 4C; S3). This result was consistent between runs (Fig. S3). We also inferred a bimodal distribution for posterior estimates for northern *R. catamitus* and *R. modestus*. We report the median time estimate for each demographic event, except in the case of bimodal distributions we report the median time of each peak, and the highest posterior density (HPD) interval of both peaks. Size changes were inferred for northern *R. catamitus* at 88.0 kya and 410 kya (95% HPD = 6.7–9,155 kya), *R. modestus* at 89.4 kya and 410 kya (27.5–683.1 kya), and northern *R. poecilonotus* at 66.2 kya (0.025–204.2 kya). In southern populations size changes were inferred for *R. catamitus* at 408.5 kya (295.8–514.1 kya), in *R. modestus* at 422.5 kya (338.0–528.5 kya), and in *R. poecilonotus* at 422.5 kya (77.4–985.9 kya). When we repeated the analysis with the admixed individual assigned to the southern population, we observed a bimodal distribution for both northern and southern *R. poecilonotus*, with more posterior probability placed on the younger event in the southern population, and a marked shift toward the older event in the northern population (Fig. S5).

**Figure 4:**
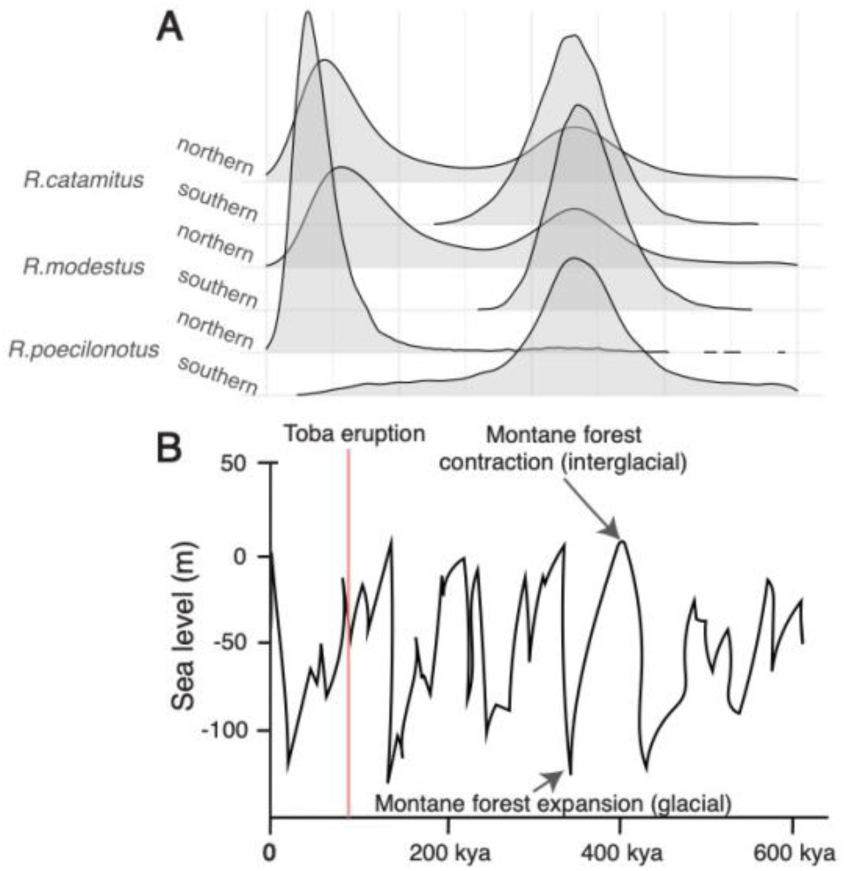
Temporal clustering of demographic events as inferred by ecoevolity v.0.3.2 (Oaks et al., 2019a) for the full-locus dataset. A) Posterior distributions of estimated demographic event times for the six populations estimated using the full-locus dataset. Times are in units of expected substitutions per site and were converted to years by dividing by the mutation rate of 1.42 x 10^-9^ substitutions per site per year (see main text). The three northern populations are clustered close to present day, with an estimated demographic event time = ∼ 80 kya. Southern populations are clustered ∼400 kya. B) Estimates of global sea levels adapted from Sarr et al., (2019) demonstrating timing of glacial cycles. During glacial periods sea levels were low, and global temperatures were cooler, thus promoting montane forest expansion. Alternatively, during interglacial periods, sea levels were higher and global temperatures were warmer, leading to refugial montane forests. The red line shows the timing of the Toba eruption ∼74 kya. Note that time in A and B moves from 600 kya on the right, to present day on the left.

The analysis inferred expansion in all populations except southern *R. poecilonotus* (Table 4). In all three species estimates of current effective population sizes encompassed far less uncertainty in the southern populations compared with the northern populations. In all cases the magnitude of size change was much greater in southern compared to northern populations.

In the exonic dataset, four demographic events received the highest posterior probability (PP = 0.42; Fig. S4B). The analysis inferred three clusters of events, the most recent shared by northern *R. catamitus* (112.6 kya, 64.5–104.8 kya) and northern *R. poecilonotus* (119.7 kya; 84.5–140.8 kya), an older event in southern *R. catamitus* (253.5 kya, 190.1–395.2 kya), and the oldest group of events including northern and southern *R. modestus* (387.3 kya, 345.1–429.6 kya) and southern *R. poecilonotus* (366.2 kya, 84.5–704.2 kya; Fig. 4D). In the analysis with the admixed individual assigned to the southern population, the northern population showed a multimodal posterior distribution while the southern population exhibited little change (Data Accessibility). We again inferred expansion in all populations, except southern *R. poecilonotus*, and the magnitude of change was greater in southern populations than northern populations, except in *R. poecilonotus* (Table 4).

### Simulated datasets

Simulations revealed that in both datasets ecoevolity struggled to accurately estimate the event times (Fig. S8). Between the two datasets, the full-locus data had a higher proportion of estimates for which the 95% credible interval contained the true value *p*(*t* ∈ CI), but the difference was minor (0.847 vs. 0.835; Fig. S8). This pattern also held when comparing the *p*(*t* ∈ CI) for each dataset between most population comparisons (e.g. *R. catamitus* north exonic and full-locus; see Data Accessibility). When comparing the true number of events to the estimated number of events, in both datasets, ecoevolity tended to overestimate the number of events, but the proportion of estimates for which the 95% credible interval contained the true value was high in all three datasets (>0.90; Fig. S9). Finally, our simulated data estimated ancestral population sizes close to the true values (Fig. S10). Although more variation is present in the descendant population estimates, the estimated values approximate the true values to a greater degree; *p*(*t* ∈ CI) = 0.823, 0.843, 0.833 for exonic, full-locus, and full-double respectively, giving us confidence in the inferred directionality of size change (Fig. S10). We found that increasing sample sizes made little if any difference on the accuracy and precision of event time estimation (Fig. S8). First, the full-double simulations did not outperform the full-locus simulations, and in many population specific comparisons actually had a lower proportion of estimates for which the 95% credible interval contained the true value. Likewise, we compared accuracy of demographic event time estimates for populations with larger and smaller population sizes and observed only marginal differences. For example in the simulations based on the full-locus dataset, *R. catamitus* south included 16 alleles and *p*(*t* ∈ CI) = 0.837, while *R. poecilonotus* south only included four alleles and *p*(*t* ∈ CI) = 0.773. In the full-doubled simulations, *R. catamitus* south had 32 alleles and *p*(*t* ∈ CI) = 0.833, compared with *R. poecilonotus* south with eight alleles with *p*(*t* ∈ CI) = 0.823 (Data Accessibility).

## DISCUSSION

### Demographic impacts of the Toba eruption on northern Sumatran *Rhacophorus*

Our results support a measurable demographic effect on northern populations attributable to the Toba eruption (Figs. 3,4,S4; Tables 3,4). Both δaδi and ecoevolity supported size change models for the northern populations of all three species, however, for *R. catamitus* and *R. modestus*, δaδi supported weak contraction while ecoevolity supported weak expansion. This conflict could simply be explained by differences in the analyses or programs, or low power in our data to estimate size changes during specific windows of time. For example, ecoevolity may have more information about older events because more coalescent events have accumulated since the size change occurred, while having less accuracy to estimate the magnitude of change at more recent time scales. Alternatively, both our δaδi model and ecoevolity assumed a single instantaneous size change event, which is clearly unrealistic considering the complexity of the geological history of the region.

From a biological perspective, this conflict could be explained by the occurrence of two size change events in these species, one at the time of the southern population expansion 300– 400 kya, and the other occurring around the time of the Toba eruption ∼80 kya (Fig. 2), thus resulting in the analysis detecting only a weak size change signal (the more recent event overriding the signal of the older event). In *R. poecilonotus* we inferred minor contraction using δaδi and minor expansion using ecoevolity but did not recover the bimodal distribution of the other two species in ecoevolity (Fig. 2), although when we reassigned the admixed individual from northern to southern, both populations exhibited weak bimodal distributions with more posterior probability placed on the older size change event (Fig. S5). Non-congruent responses to Pleistocene events between *R. catamitus* and *R. modestus* and *R. poecilonotus* may also be partly explained by their differing evolutionary histories (O’Connell et al., 2018a), or reflect yet unidentified differences in biological traits (Papadopoulou & Knowles, 2016; Zamudio et al., 2016), perhaps driven by local adaptation to different parts of Sumatra in the early stages of divergence (Losos & Ricklefs, 2009).

Although our data does not allow us to directly assess the impact of the Toba eruption on local extinction, past studies have shown that the eruption probably resulted in few extinctions (Louys 2007; Louys & Meijaard, 2010), and studies on the impacts of Holocene eruptions observed high survivorship and rapid recolonization in small-bodied animals, including amphibians (Burt, 1961; Carson et al., 1990; Crisafulli et al., 2015). Alternatively, our data concord well with past studies supporting a bottleneck that reduced local genetic diversity (Ambrose, 1996; Rampino & Ambrose, 2000; Rampino & Self, 1993a, 1993b; Nater et al., 2011; 2017). Several studies, including this one, have attributed shifts in effective population size or pulses in diversification to the Toba eruption (Louys et al., 2004; Nater et al., 2011; 2017). Nonetheless, Toba was only one of many eruptions on Sumatra during the Quarternary that impacted diversification and demography of resident taxa (Oppenheimer, 2002), and the increase of genomic sequence data for Sumatran taxa may unlock opportunities to test the effects of multiple eruptions on diversification.

### Demographic impacts of forest expansion

As expected, we inferred that southern populations exhibited higher rates of genetic diversity than northern populations (Figs. 2;S3). We expect that as more island-wide (and genome-wide) datasets are generated, disparities between the genetic diversity of northern and southern populations will be commonly identified. Although we expected to find these differences in genetic diversity, we did not expect to recover a strong signal of size change in the three southern populations much earlier than the Toba eruption (∼400 kya; Figs. 4;S4), and no demographic signal of the Toba eruption in these southern populations (Fig. 4). This calls into question studies that posit global as compared to local effects of the Toba eruption, supporting the findings of Oppenheimer (2002) and others (i.e. Yost et al., 2018). We even found that this earlier size change event may have also impacted northern *R. catamitus* and *R. modestus* (and *R. poecilonotus* depending on the assignment of the admixed individual).

We hypothesize that two processes could have driven this older size change event. First, many other eruptions occurred during the Pliocene across Sumatra and in west Java that could have driven demographic shifts in these populations (Global Volcanism Program, 2013). However, northern populations would likely not show a signal of size change at this older event if it was driven primarily by an eruption in southern Sumatra or west Java, as most examples of the genetic consequences of eruptions posit localized effects (Carson et al., 1990; Brown et al., 2000; Gubitz et al., 2005; Bloor et al., 2008).

The second hypothesis we propose is that montane forest expansion during glacial cycles every 100 kya. As montane forests expanded down-slope during glacial periods (Newsome & Flenley, 1988; Stuijts et al., 1984, 1988; Cannon, 2012), montane-forest species could disperse between previously isolated mountains, and likely saw increases in effective population sizes. Further, our SDMs supported expansion of *Rhacophorus* distributions during glacial cycles (Fig. 1B), which closely matched the montane forest distributions modelled by Cannon et al. (2009). In particular, we propose that the glacial period between ∼300–400 kya (Fig. 3B) precipitated a large size change event as montane forests expanded down-slope (Fig. 1B). This hypothesis is broadly supported by the 95% HPD of demographic size change for southern populations across both datasets (∼300–500 kya).

### Future directions

Our study highlights the difficulties of unravelling diversification histories in a region subjected to complex historical processes (Lohman et al., 2011; Frantz et al., 2013; Leaché et al., 2020). One limitation of current models is that they are restricted to testing fairly simple demographic histories (Oaks et al., 2019c; Terhorst & Song, 2015), which approximate diversification processes of taxa subjected to complex events such as glacial cycles or repeated volcanic events (Lohman et al., 2011). For example, we inferred bimodal distributions in several populations, suggesting either: (1) low support for two demographic events, (2) that our data had little power to distinguish the timing of events in these populations, or (3) these populations experienced two population size changes in the past, and because ecoevolity only models one event, the posterior is ‘split’ between them. More work is needed to understand how these classes of simple models perform in geologically complex regions.

Future studies could also further investigate the effects of different datasets on population genomic analyses. For example, using our subsampling approach, we found consistent estimates of population genetic parameters for the two target capture and ddRADseq datasets (Fig. S3). Yet it remains unclear why our ddRADseq dataset failed to converge in ecoevolity, especially as this program was developed using RADseq sequence data (Oaks et al. 2019a,c). Future studies could use simulations to further compare target capture and ddRADseq datasets for population genomic analyses and identify if differences in demographic inference results could be due to higher information content of the target capture loci or due to the types of SNPs generated from protein-coding genes and random ddRADseq-derived SNPs. See the Supporting Information for more discussion on this subject.

Finally, we inferred large differences between the full-locus and exonic datasets in posterior estimates in ecoevolity (Fig. S4) and in some cases between population genetic parameter estimates (Fig. S3). Simulations in pycoevolity of both datasets suggested that differences observed between estimates from the empirical datasets were not due to differences in the number of sites or missing data patterns, but rather may be due to differences in signal from the sampled character patterns. These signals could also be driven by biological factors such as differential selection on exons compared with flanking regions, or data collection patterns such as paralogy and acquisition biases. Further work could use simulation or empirical approaches to investigate how these different data types perform under various evolutionary scenarios such as demographic events occurring at recent or ancient time scales.

## ACKNOWLEDGEMENTS

We thank four anonymous reviewers for their critical feedback on this manuscript. We thank the Ministry of Research and Technology of the Republic of Indonesia, for granting permits for this project, and representatives of LIPI for housing specimens and facilitating permits. We thank members of the field team for helping to collect voucher specimens. We are grateful to Dr. Jimmy McGuire (MVZ), and Drs. Stefan Hertwig and Manuel Schweizer (NMB) for tissue loans. We thank Jose Maldonado for generating RNAseq data, Dan Portik for sage advice on wet lab work, Utpal Smart for help at the bench, and everyone at Smithsonian Data Science Lab (plus Vanessa González at GGI) for bioinformatics support. Finally, we thank the members of the Bell lab at the NMNH for their valuable feedback on early drafts of this manuscript. A National Science Foundation Doctoral Dissertation Improvement Grant to MKF and KAO [DEB-1701721], a Division of Environmental Biology grant to ENS and M.B. Harvey [DEB-1146324], and a Global Genome Initiative Peter Buck Postdoctoral Fellowship to KAO funded this work.

## DATA ACCESSIBILITY

Tissues used to generate transcriptome data were both loaned from the Museum of Vertebrate Zoology (MVZ), University of California Berkeley by Jimmie McGuire (JAM). Double-digest RADseq data from O’Connell et al., (2018a) can be found on the SRA database under SAMN05426771–SAMN05426803, and SAMN08437163–SAMN08437177. Demultiplexed raw data from the target capture data can be found at SRA XXX [deposited after acceptance]. Workflow for transcriptome-based capture probe design can be found at: https://github.com/kyleaoconnell22/Exome_Capture_Probe_Design.git Workflow for filtering locus alignments post SECAPR/PHYLUCE and for calling a consensus sequence from phased alignments is found at: https://github.com/kyleaoconnell22/TargetCaptureAnalyses. Workflow for subsampling populations for parameter estimation https://github.com/kyleaoconnell22/Pop_gen_estimation_with_random_subsampling. Alignment files and ecoevolity configuration files and figure outputs are available at Figshare doi XXX [deposited after acceptance]. Files used to generated species distribution models are available at: https://github.com/kshaney/Sumatra_ConBiogeo.

## AUTHOR CONTRIBUTION

E.N.S., A.H., N.K., K.J.S. and K.A.O. conducted fieldwork; K.A.O. and M.F.K. conducted laboratory work; K.A.O., M.F.K. and E.N.S. procured funding, K.A.O., J.R.O. and K.J.S. ran analyses; K.A.O. led the writing with input from all authors, and all authors approved of the final version of the manuscript.

## Supporting Information

### INTRODUCTION

#### A note on taxonomy of Rhacophoridae

Jiang et al., (2019) used 972 bp of mitochondrial sequence from 55 species and morphological data from the literature to revise the taxonomy of the genus *Rhacophorus*. In their paper they used phylogenetic analyses of the mitochondrial data to infer three well-supported clades within *Rhacophorus* which had been recovered in past studies (e.g. O’Connell et al., 2018a). They divided *Rhacophorus* into three genera, applying the name *Rhacophorus s.s.* to the group of species encompassing most Sundaland taxa (including most of the Sumatran and Javan species). The clade that includes most of the taxa from Borneo (as well as *R. cyanopunctatus* from across Sundaland including on Sumatra), they assigned to the generic name *Leptomantis*. Even though *Leptomantis* was synonymized with *Rhacophorus s.l.* by Harvey et al., (2002) because it was non-monophyletic with respect to *Philautus* and the characters used to define it applied to members of *Rhacophorus s.s*. Finally, the clade including mostly East Asian taxa Jiang et al., (2019) assigned to the new genus *Zhangixalus*. Although several past studies have consistently recovered these three clades, the study by Jiang et al., (2019) contains a number of shortcomings that should be rectified before their new taxonomy is adopted.

First, they failed to sample all available sequences on Genbank (at least 68 at the time of O’Connell et al. 2018b), instead only including 55 of the 95 species described from the genus (amphibiaweb.org). Incomplete taxon sampling is known to influence phylogenetic estimation (Zwickl & Hillis, 2002), and this may have serious implications when revising whole genera based on limited sampling. For example, the study failed to include any sequences of *R. cyanopunctatus*, although sequences from multiple landmasses are available on Genbank and were sampled by past studies (e.g. O’Connell et al. 2018a,b). In addition, with the availability of an abundance of nuclear data on Genbank (e.g. Chan et al. 2018; O’Connell et al. 2018a,b), it seems inappropriate to only sample mitochondrial data, particularly when assigning clades to genera. The primary issue for our study is that some taxa from Sumatra have been placed in the phylogeny in different locations in different studies. For example, Jiang et al., (2019) placed the *R. achantharrhena* clade as part of *Zhangixalus* with strong support, which was also recovered by Li et al., (2012) using only mitochondrial DNA. Chan et al., (2018) recovered a similar topology using nuclear and mitochondrial DNA, whereas O’Connell et al. (2018a,b) used nuclear and mitochondrial data to place the *R. achantharrhena* clade as sister to both the clade referred to as *Leptomantis* and that referred to as *Zhangixalus*. Jiang et al. (2019) admit that the placement of the *R. achantharrhena* clade remains uncertain, but assign these species to their new genus, even though they do exhibit any of the characters used to define the new genus (see Discussion of Jiang et al. 2019). Finally, Chen et al. (2020) does resolve the relationships of these three clades using genomic sequence data, but because their study was focused more on familial relationships, their sampling within *Rhacophorus s.l.* was limited. For example, their only representative of *Leptomantis* was *R. cyanopunctatus*, which was lacking from the Jiang et al. (2019) study. Due to the uncertainty of placing some of the Sumatran species in the revised taxonomic arrangement, we advocate for not adopting this new taxonomy until more comprehensive phylogenetic sampling is undertaken, specimens are examined to confirm the characters for each clade (they only used characters from the literature), and perhaps the phylogeny is better resolved using phylogenomic analyses rather than a few mitochondrial and nuclear genes.

### MATERIALS AND METHODS

#### Transcriptomic data generation and analysis

Two species of *Rhacophorus* were chosen for transcriptome sequencing: *R. modestus* (MVZ 272194) from Sumatra, and *R. monticola* (JAM 14420) from Sulawesi (see Data Accessibility for museum information). These species were chosen to target at least one species from each of the two primary clades within *Rhacophorus* (O’Connell et al., 2018b). Whole RNA was extracted from liver samples flash frozen in RNA-later. RNA was quantified using the Agilent 2100 Bioanalyzer (Aligent Technologies, Santa Clara, CA, USA) with an RNA chip kit. Extracted RNA for each sample was sent to the Brigham Young University DNA Sequencing Center (dnasc.byu.edu) for library preparation and sequencing on a single lane of the Illumina® HiSeq 2500.

Raw reads were cleaned and *de novo* assembled into contigs using TRINITY v.2.1.1 (Grabherr et al., 2011) under default settings. We identified orthologous exons between the two species following Yang and Smith (2014). Briefly, putative coding regions were identified with TransDecoder v.2.10 (Haas et al., 2013) using the *Nanorana parkeri* genome. Redundant reads were removed with cd-hit-est v.4.64 (Li & Godzik, 2006). Bastn v.2.7.1 (Altschul et al., 1990) was used to remove contigs from each individual that were not present in both individuals. Markov clustering was performed using MCL v.1.30 (Enright et al., 2002), which clusters sequences based on the fraction and number of blast hits. A phylogenetic tree was inferred for each homolog, and finally, the data were filtered to 1-to-1 orthologs and aligned using MAFFT v.7.302 (Katoh et al., 2002). We identified 2828 orthologous transcripts and used these to design target loci for capture.

#### Target capture probe design

Target sequences were designed using a custom pipeline (See Data Accessibility in main text). We used blasn v.2.7.1 (Altschul et al., 1990) to blast sequences from MVZ 272194 against JAM 1420 to identify portions of sequence alignments without ambiguous sites or gaps. We retained only completely overlapping regions, then trimmed sequences to the first 1000 bp using fastx_trimmer v.0.0.13 (http://hannonlab.cshl.edu/fastx_toolkit/). We retained alignments with GC content between 40–70 percent, sequence divergence between 1–15 percent, and sequence length between 430–1000 bp. We used RepeatMasker v.4.09 (Smit et al., 2013) to remove any transcripts with repetitive content. Probes with high or low GC, or that are too divergent from the target species can perform poorly during hybridization (Bi et al., 2012; Bragg et al., 2016; Portik et al., 2016). Probes for the target sequences were synthesized by myBaits® (Arbor Biosciences, Ann Arbor, MI) using 120 bp probes with 2x tiling. The final probe set resulted in 19,950 baits targeting 955 transcript markers from each of the two species.

#### Genomic library preparation

Genomic DNA was extracted from liver and muscle tissue stored in Cell Lysis Buffer from 69 individuals (Table SI) using a standard salt-extraction method (Sambrook & Russell 2001) followed by an additional cleanup with Sera-Mag Speedbeads (Rohland & Reich, 2012). Concentrations were checked on a QUBIT® 2.0 Fluorometer (Life Technologies, Grand Island, NY, USA), and size distributions were checked on a 1% Agarose gel using 1–5 μl of DNA. We fragmented 300–800 ng of DNA per sample using 2 μl of NEB dsDNA Fragmentase (New England Biolabs, Ipswich, MA) for 20–25 minutes at 37°C and prepared genomic libraries using the Kapa Hyper Prep Kit (Kapa Biosystems, Wilmington, MA) using half the manufacture’s recommended reaction volumes. For our first three library groups (all species except *R. modestus*, *R. poecilonotus*, and *R. achantharrhena*), we did not size select between adapter ligations and PCR amplification. We conducted two PCR reactions of 7–8 amplification cycles under the following conditions: 98◦C for 45 sec; 7–8 cycles of 98◦C for 30 sec, annealing temperature of 60◦C for 30 sec, 72◦C for 30 sec, final extension of 72◦C for 1 min, and final rest at 12◦C. We pooled individuals into three capture groups of 7–8 individuals. Because the size distribution of these three libraries was very wide, we decided to size select our remaining libraries using the Blue Pippin electrophoresis platform (Sage Science, Beverly, MA, USA) for fragments between 450–750 bp. Thus, with the remaining 8 capture groups, we pooled 5–7 individuals following adapter ligation, size selected the pool, then PCR amplified pools in 4–6 reactions of 8 cycles. We checked the distribution of each pooled library group on the Agilent 2100 Bioanalyzer.

#### Hybridization reactions

Starting with 1 μg of DNA from each pooled library group, we followed Portik et al., (2016) to utilize two variations of blocking oligos, including the myBaits® supplied blocking oligos, as well as IDT xGen blocking oligos (Integrated DNA Technologies). When using the xGen blockers, we added 2.5 μl of the B1, 1 μl of the B2, and 2 μl of the xGen blockers. We incubated hybridization reactions for 24 hours at 65◦C. We amplified hybridized DNA in three reactions of 14 PCR cycles, with the exception of the three groups that were not size selected (Table S1) which we amplified for 30–36 cycles in three reactions because initial concentrations at 14 cycles were too low. We confirmed the distribution of each capture reaction on the Agilent 2100 Bioanalyzer, and quantified total library concentration using the QUBIT. Finally, we pooled capture libraries into two sequencing groups (Table S1) which were sequenced on two lanes of (150 bp PE; lane 1 was shared with another project, see Table S1) of the Illumina® X10 at Medgenome (medgenome.com).

#### Genomic data processing

We demultiplexed barcoded samples using FASTX barcode splitter v.0.0.13 (http://hannonlab.cshl.edu/fastx_toolkit/), and conducted remaining data processing using the Python pipeline SECAPR v.1.14 (Andermann et al., 2018), which is an extension of the Phyluce pipeline (Faircloth, 2015). Our SECAPR workflow included the following steps: (1) we cleaned raw reads using TRIMMOMATIC v.0.36 (Bolger et al., 2014), hard trimming the first 10 bp and last 13 bp from each read to remove adapter sequence, and setting the other parameters as: simpleClipThreshold = 5, palindromeClipThreshold = 20, and seedMismatch = 5. We checked the quality of each sample using FastQC, and only proceeded after all samples passed QC tests. (2) We assembled cleaned reads into contigs using TRINITY v.2.1.1 (Grabherr et al., 2011). We updated SECAPR to v.1.15 and separated our target loci by species (*R. modestus* and *R. monticola*) but after finding little difference between the two references, we used *R. modestus* target sequence for all subsequent analyses. (3) We matched assembled contigs to the target sequences for each of the three focal species separately with min-coverage = 60, min-identity = 60, and used the keep-duplicates and keep-paralogs flags. We aligned the recovered markers for each species using MAFFT v.7.407 (Katoh et al., 2002). (4) We conducted reference based assembly using the reference_assembly script for each species by mapping our cleaned reads to the species-specific reference (reference type = alignment consensus). (5) We phased alleles (phase_alleles, min coverage 6x per allele) and again aligned using MAFFT (no trim and allow ambiguous). We further filtered by removing sequences with more than 80% missing data at a locus and trimmed using GBLOCKS v.0.91b (-b2=.1 -b3=8 -b4=10 -b5=a -t=d; Castresana, 2000). We then removed trimmed sequences shorter than 50 bp, and sequences with more than 45% missing data. We refer to this data set as ‘full-locus’. All scripts used in filtering are available online (see Data Availability).

In order to optimize our alignments at the phylogenetic level (across the genus) and also to evaluate the utility of exonic sequence for population genomic analyses, we generated a second data set that we refer to as the ‘exonic’ data set. Rather than assembling reads then matching the assembled contigs to a reference sequence, we mapped our cleaned reads directly to our target sequences, then phased, aligned, and filtered these alignments as described above.

#### Details on ecoevolity parameter variations

We applied a Dirichlet process with a concentration parameter values fixed at 2.62 on the event model prior (estimate = false), which corresponds to at least half the prior probability being placed on 3.5 demographic events, although we also tested prior values of 12 (5 events) and 20 (5.3 events). We applied an exponential distribution to the event time prior with a rate of 1000 (corresponding to a mutation-rate converted value in years of ∼500 Kya), but also tested values of 6000 (∼95 Kya), 800 (800 kya), 500 (∼1.4 Ma), and 100 (∼5.6 Ma). Under global comparison settings, we set genotypes = haploid, and set constant sites removed = false, and equal population sizes = false. For the prior on the current population size, we used a theta = 0.004 (calculated from the full-locus data set from each species) to place a prior value of 0.001 with a gamma distribution (shape = 3.5, scale = 0.001). We visualized our prior distributions using (https://github.com/ivanprates/2018_Anolis_EcolEvol/tree/master/scripts), ensuring that the gamma distribution encompassed our starting value. For the root relative population size, we used our δaδi results to apply population-specific exponential distributions. On the northern populations of *R. catamitus* and *R. modestus* we applied an exponential distribution with a mean = 2, suggesting that the ancestral population was twice as large as the current population (contraction scenario), and on the southern populations applied an exponential distribution with a mean = 0.5, such that the current population is twice the size of the ancestral population (expansion). In *R. poecilonotus* we applied the inverse parameters. In all cases we allowed the parameter to be estimated. We also tested the inverse of these size priors to test their sensitivity on analyses but observed no differences in posterior estimates.

### RESULTS

#### Target capture efficiency at the generic level

We filtered out samples that captured fewer than 20% of loci (<190 loci), leaving 45 samples for downstream analyses (Table S1). Our filtered sampling included seven nominal species from across the *Rhacophorus* phylogeny including *R. achantharrhena* (n = 3; mean loci captured in exonic data set = 592), *R. bengkuluensis* (n = 1; 917 loci), *R. nigropalmatus* (n = 1; 898 loci), *R. reinwardtii* (n = 1; 877 loci)*, R. catamitus* (n = 13; 878 loci), *R. modestus* (n = 15; 887 loci), *R. poecilonotus* (n = 10; 819 loci), and one species from a closely related genus, *Feihyla kajau* (n = 1; 699 loci). Generally, our probe set worked well across the genus and with the outgroup sample, recovering 950/951 target loci across a minimum of three samples. To illustrate, we recovered an average of 887 loci (765–950) for *R. modestus* the species from which transcriptome-based probes were designed, while the outgroup *F. kajau* retained 699 loci across ∼36 Ma divergence (O’Connell et al. 2018b).

#### Phylogenetic results

Our ML phylogeny of all species matched closely with that of O’Connell et al., (2018b) based on multi-locus Sanger sequence data (Fig. S2). We inferred *R. achantharrhena* as sister to all other Sumatran and Javan *Rhacophorus*. We recovered *R. nigropalmatus* as sister to *R. reinwardtii*, and these two species were sister to the clade that included the focal Sumatran species. Within the focal Sumatran clade, we inferred *R. bengkuluensis* from Java as sister to *R. catamitus*, and these species were sister to *R. modestus* and *R. poecilonotus*. Only the node beween *R. nigropalmatus* and *R. reinwardtii* had less than 100% bootstrap support.

### Discussion

#### Comparing target capture and ddRADseq data for population genomic studies

Studies comparing target-capture data to ddRADseq data for phylogenomic inference generally find that the two data types perform similarly for estimating phylogenies, although performance may differ at nodes of various depths (Leaché et al., 2015; Collins & Hrbek, 2015; Harvey et al., 2016; Manthey et al., 2016). Yet few studies have evaluated possible differences between data types for studying population level processes. A number of factors may affect population genomic analyses such as parameter estimates and demographic modelling, particularly those that depend on the allele frequency spectrum (Harvey et al., 2016). In particular, levels of missing data and information content per locus (Collins & Hrbek, 2015) may affect both model selection and parameter estimation, but this has not been well quantified by direct side-by-side studies. Because our sampling was not matched at the individual level, it is difficult to parse out the possible reasons for observed discrepancies between datasets across analyses. By taking a resampling approach for genetic diversity parameter estimates, we found that the three datasets inferred similar parameter estimates, except in a few cases with the smallest sample sizes which may have been subject to the most between-sample variation (Fig. S3). Nonetheless, we were unable to take a similar approach with the modelling analyses, and with both δaδi and ecoevolity it remains unclear why the target capture and ddRAD data performed differently. Ultimately, we attribute differences to the higher information content of the target capture loci, but it also may be due to the types of SNPs generated from protein-coding genes and random ddRADseq-derived SNPs. Future studies could use simulations to further compare target capture and ddRAD datasets for population genomic analyses.

---

**Figure S1:**
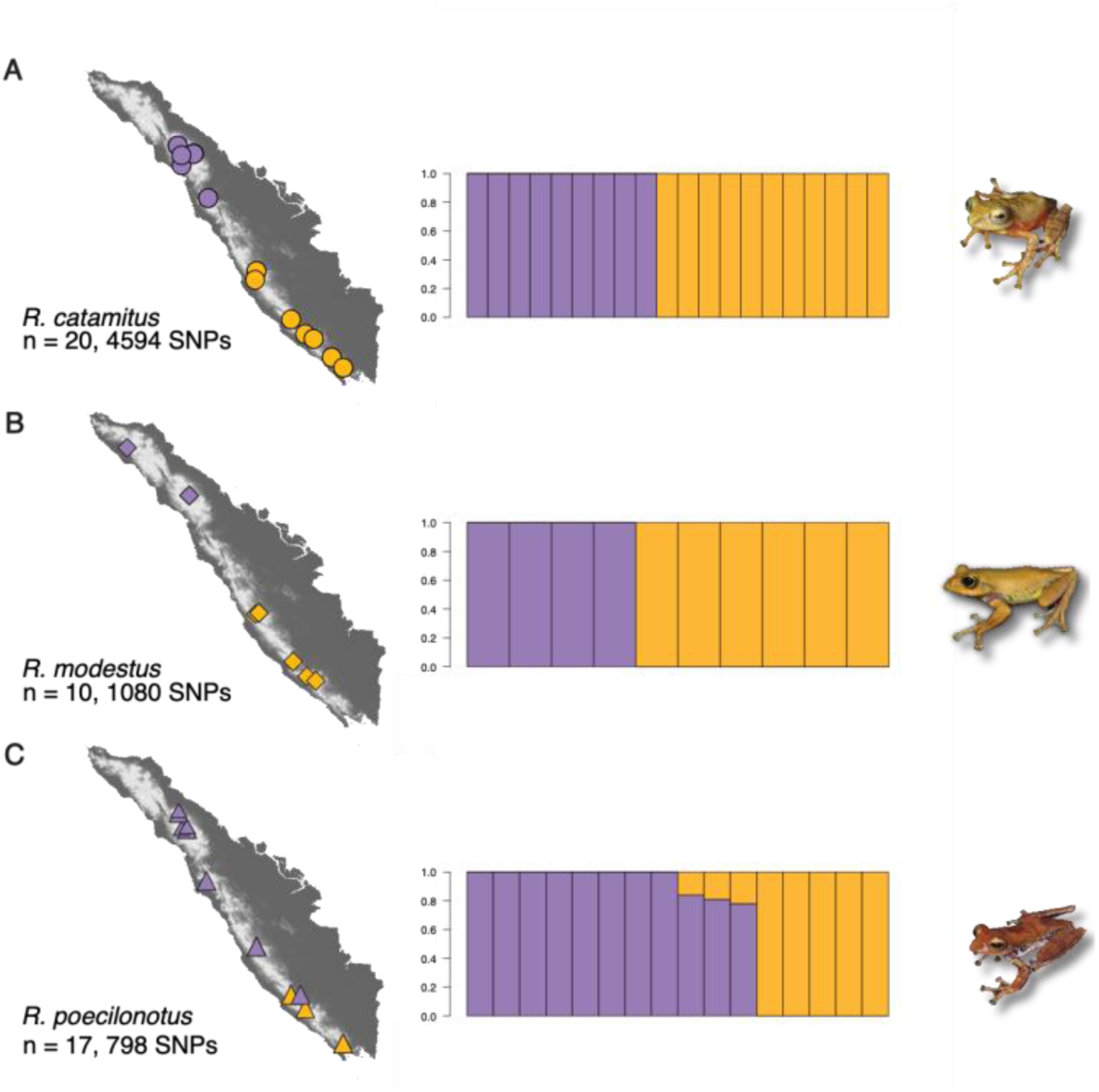
Sampling for ddRADseq datasets from O’Connell et al (2018a). Maps demonstrate the geographic location of each sample, and are accompanied by ancestry proportions estimated in ADMIXTURE v.1.3.0 (Alexander et al., 2009). Individuals from northern populations are coloured purple, southern populations are coloured orange. Maps and admixture proportion plots correspond to A) *R. catamitus*, B) *R. modestus*, C) *R. poecilonotus*.

**Figure S2:**
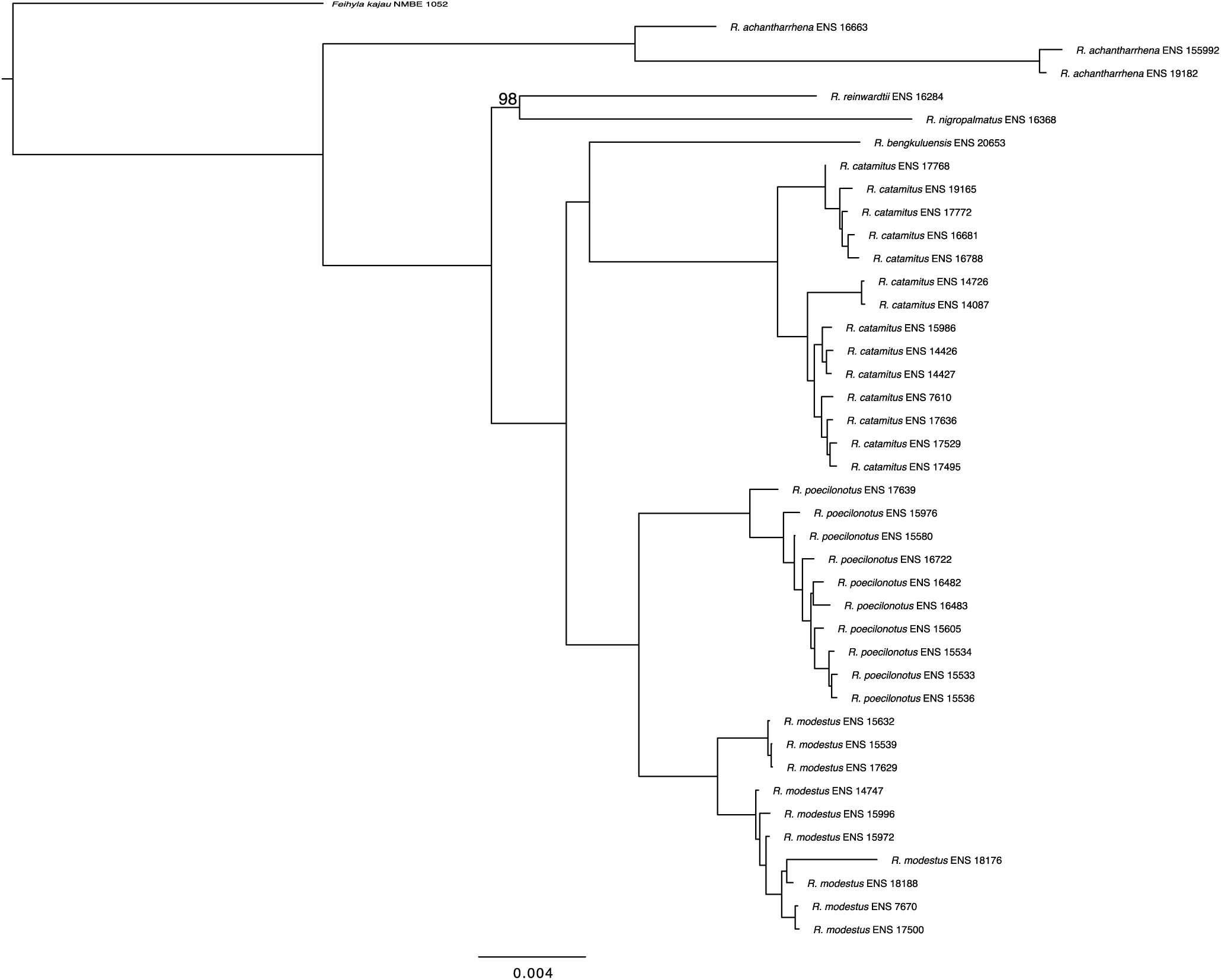
Phylogeny of consensus sequences for all samples from the capture experiment that passed filtering generated using RAxML v.8.2.11 with 100 rapid bootstrap iterations. The only node with bootstrap support <100 was the node between *R. reinwardtii* and *R. nigropalmatus*. Analysis is based on 944 loci from the exonic dataset, and recovers phylogenetic relationships matching past studies (e.g. O’Connell et al. 2018b; Chen et al., 2020).

**Figure S3:**
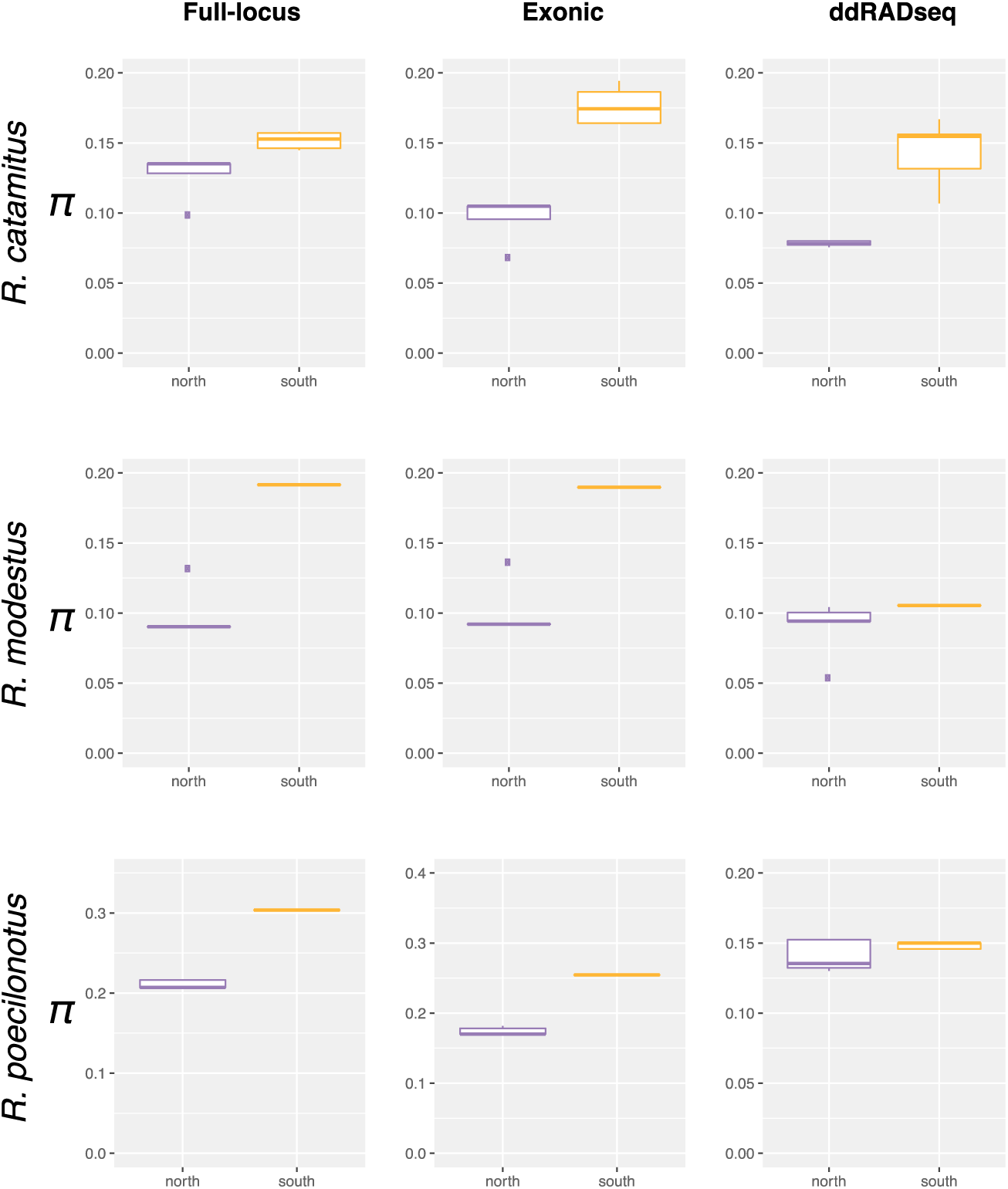
Estimates of nucleotide diversity (π) between northern and southern populations for each species and each of the three datasets after controlling for sample size discrepancies by resampling (see main text). Northern populations are coloured purple, southern populations are coloured orange. Results show general consistencies between datasets using resampling, and in all cases the southern populations have higher genetic diversity than northern populations, except in *R. poecilonotus* ddRADseq.

**Fig. S4:**
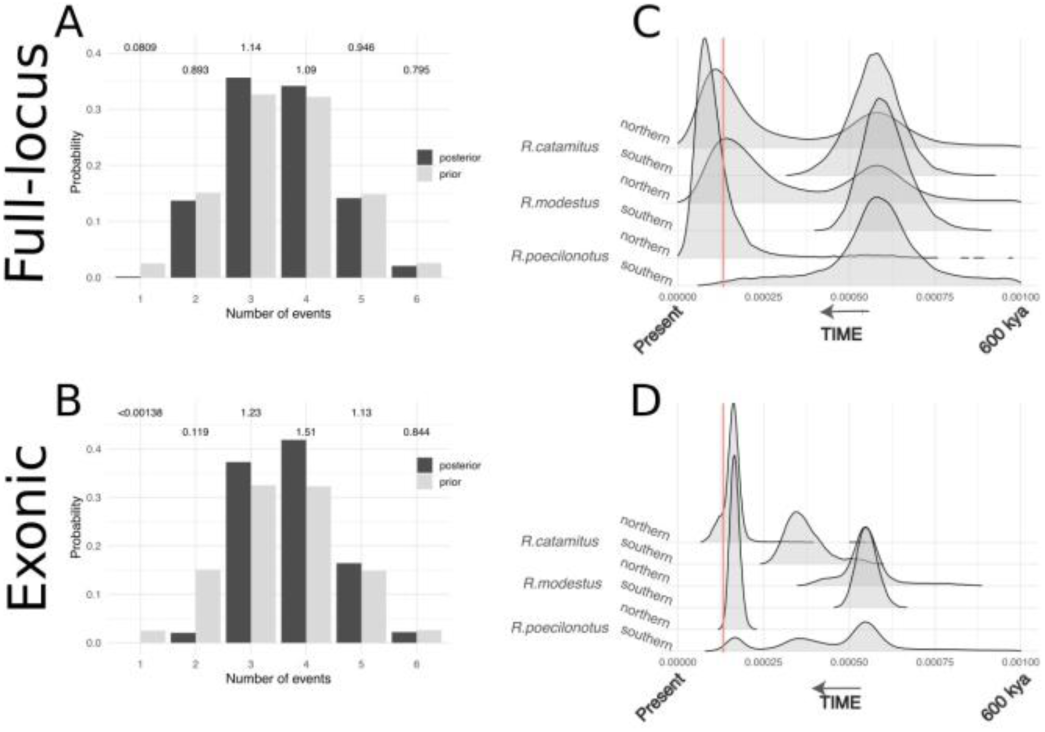
Temporal clustering of demographic events as inferred by ecoevolity v.0.3.2 showing estimates for the number of events, as well as a comparison of the timing of events when estimated using the full-locus and exonic datasets. A,B) Posterior (dark gray) and prior (light gray) probability for the number of demographic events, with the Bayes Factor above each set of bars, shown for the A) full-locus dataset, B) exonic dataset. C-D) Posterior distributions of estimated demographic event times for C) full-locus dataset, D) exonic dataset. Times are in units of expected substitutions per site and were converted to years by dividing by the mutation rate of 1.42 x 10^-9^ substitutions per site per year. The red line shows the timing of the Toba eruption.

**Fig. S5:**
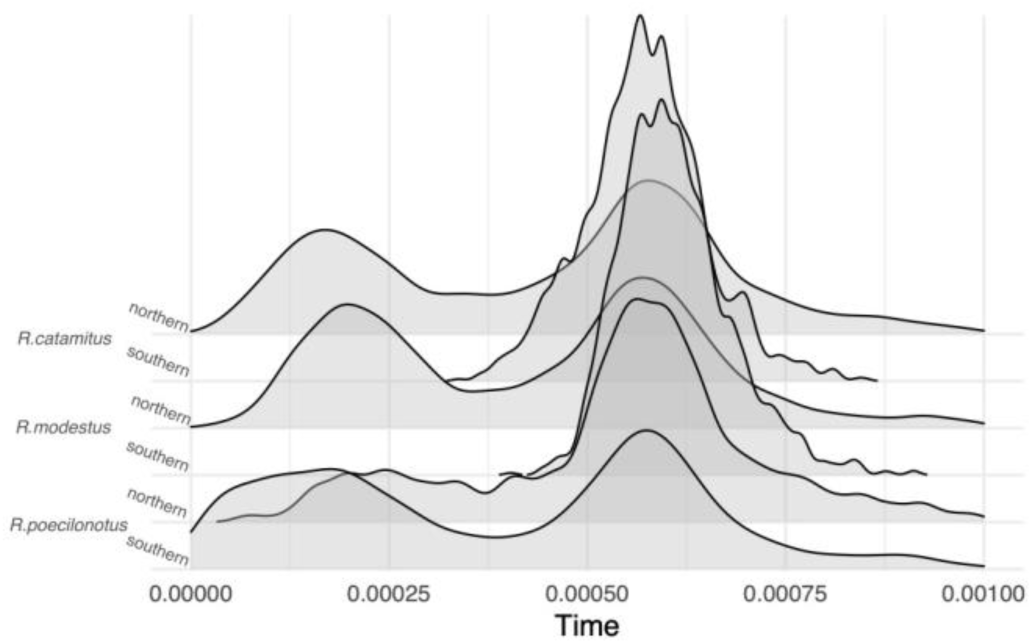
Temporal clustering of demographic events as inferred by ecoevolity v.0.3.2 for the full-locus dataset after reassigning the admixed *R. poecilonotus* individual from northern to southern. The change drives the *R. poecilonotus* northern posterior event times closer to the older demographic event, but with a bimodal distribution split between the two events. The analysis also infers a bimodal distribution for the southern *R. poecilonotus* with more of the posterior placed on the recent event. This suggests that although admixed, this individual carries more signal of the Toba eruption indicative of the northern population.

**Figure S6:**
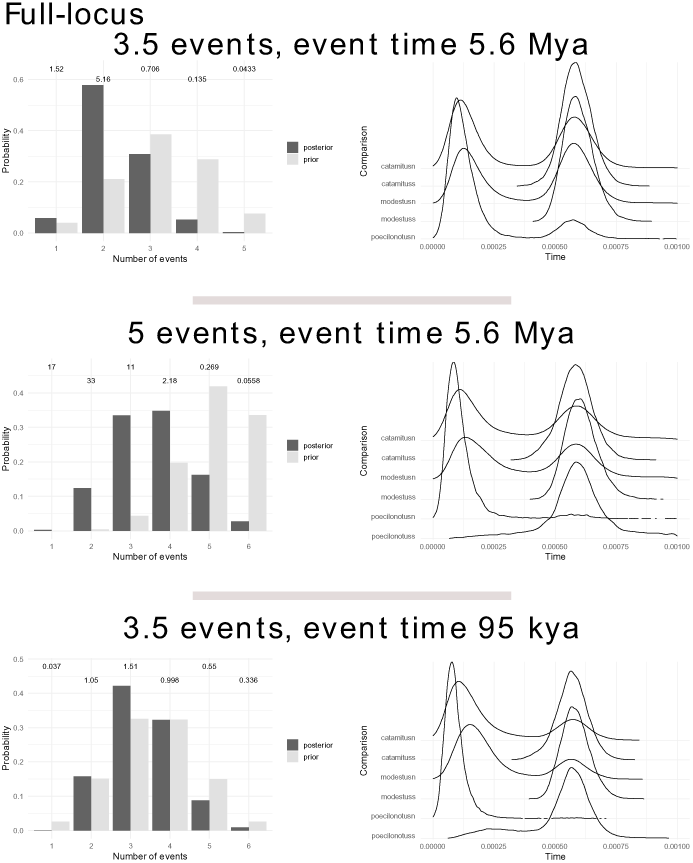
Examples of outputs for the full-locus dataset from ecoevolity when testing alternative parameter combinations. Plots show the effect of the prior on the concentration parameter on the prior and posterior support for the number of demographic events, but not the posterior distribution of event times. The number of events above each figure signifies the mean prior probability on that number of events, while the event time is the mutation-rate-converted rate of the exponential distribution on the event time prior.

**Figure S7:**
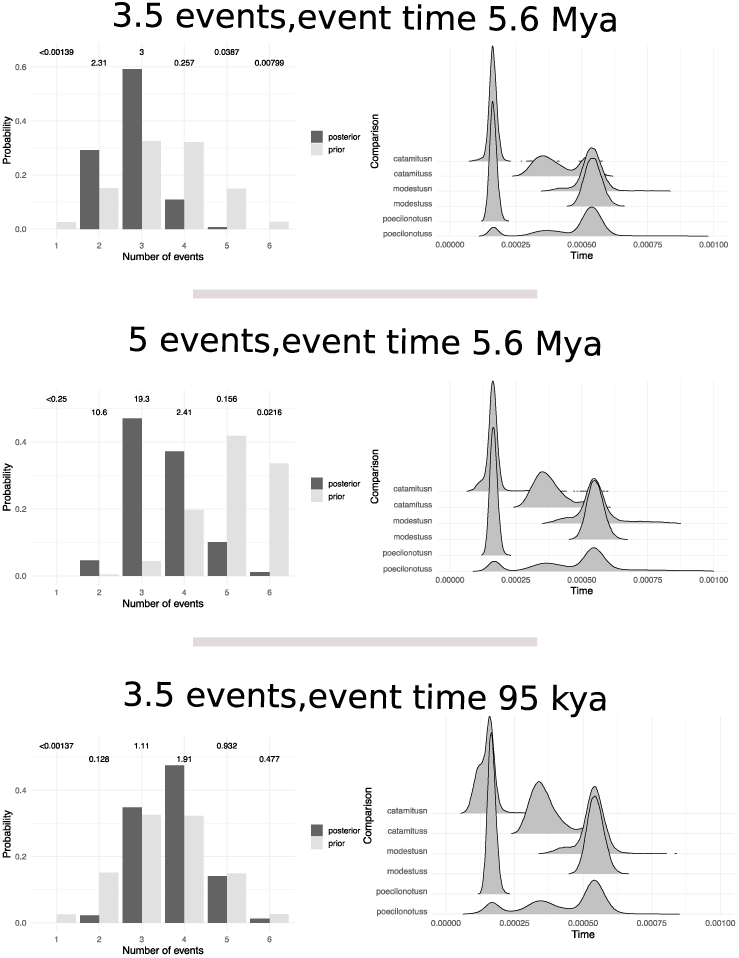
Examples of outputs for the exonic dataset from ecoevolity when testing alternative parameter combinations. Plots show the effect of the prior on the concentration parameter on the prior and posterior support for the number of demographic events, but not the posterior distribution of event times. The number of events above each figure signifies the mean prior probability on that number of events, while the event time is the mutation-rate-converted rate of the exponential distribution on the event time prior.

**Figure S8:**
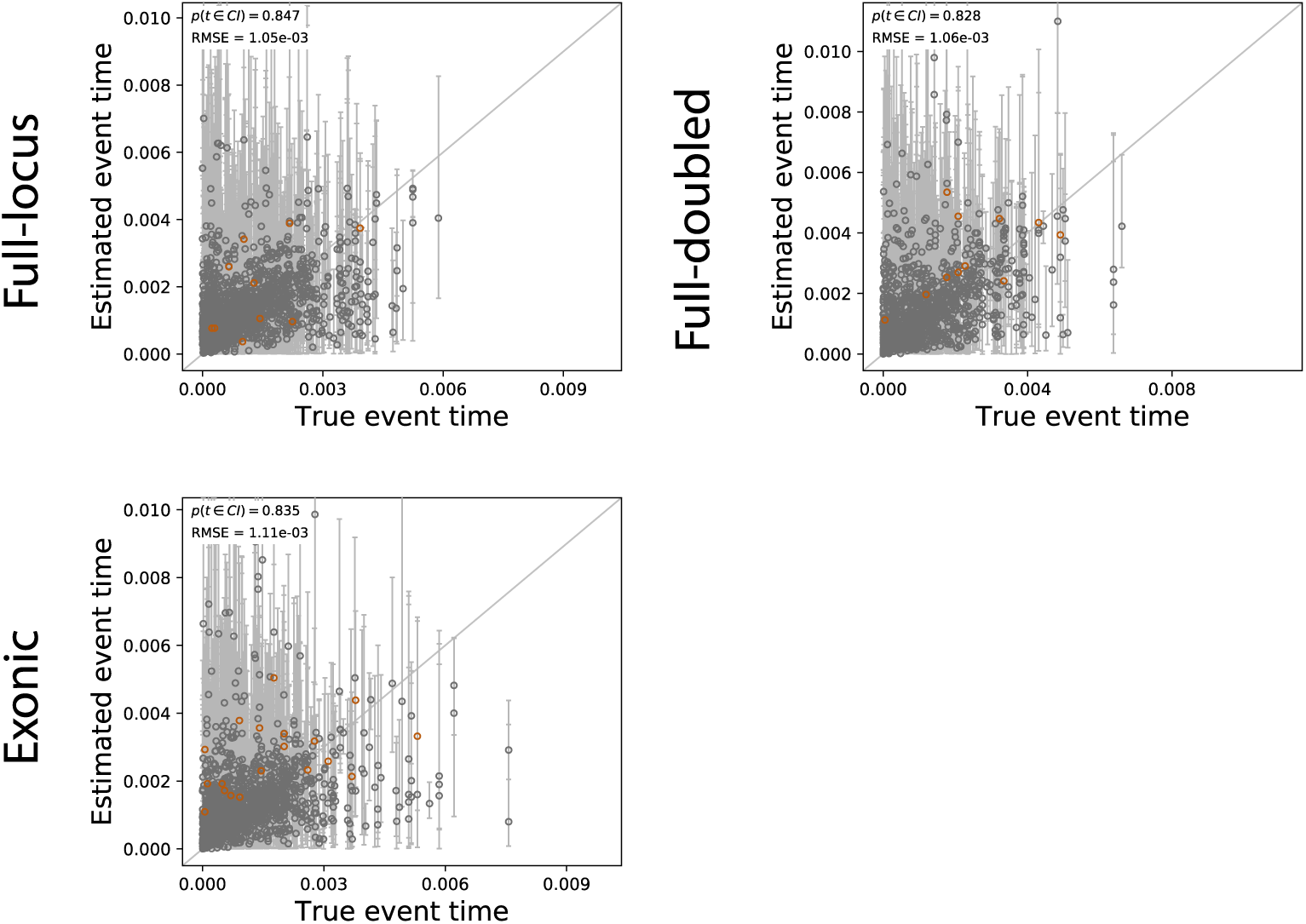
The accuracy and precision of estimates of demographic event times (in units of expected substitutions per site). Here we compare data simulated to match our empirical data for numbers of individuals, locus number, locus length, and patterns of missing data within each locus. We also simulate one dataset with double the number of individuals to better understand the effect of sample size on posterior estimates (full-doubled). The full-locus and full-double datasets are on the top row, while the exonic dataset is on the bottom row. Each plotted circle and error bars represent the posterior mean and the 95% credible interval. Estimates for which the potential scale reduction factor was greater than 1.2 (indicating poor convergence/mixing) are highlighted in orange. Each plot contains 1,800 estimates (300 simulated replicates each with six time estimates). Each plot shows the root-mean-square error (RMSE) and the proportion of estimates for which the 95% credible interval contained the true value – (*p*(*t* ∈ CI). Plots were generated using ‘pycoevolity’ v.0.2.4.

**Figure S9:**
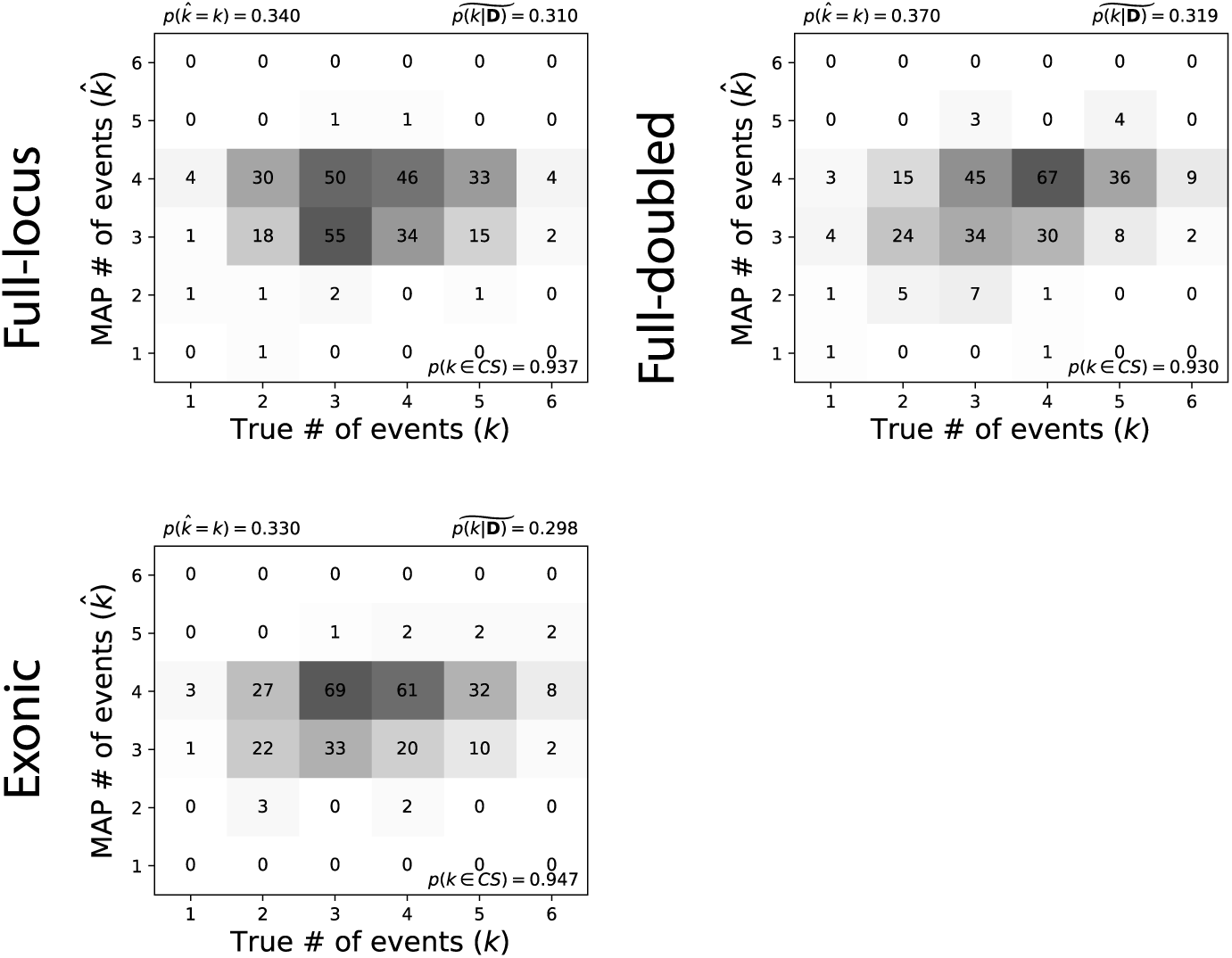
The performance of ecoevolity to estimate the true number of events using data simulated to match our empirical data, as well as one simulated dataset with twice as many samples. Each plot shows the results of the analyses of 300 simulated replicates, each with six demographic event estimates. The number of datasets that fall within the cells of the true verses estimated model is shown, and cells with more data have darker shading. The estimates are based on the model with the maximum *a posteriori* probability (MAP). For each plot, the proportion of datasets for which the MAP model matched the true model is shown in the upper left-hand corner, and the median posterior probability of the correct model across all datasets is shown in the upper right-hand corner. The proportion of estimates for which the 95% credible interval contained the true value is shown in the bottom right corner. Plots were generated using ‘pycoevolity’ v.0.2.4.

**Figure S10:**
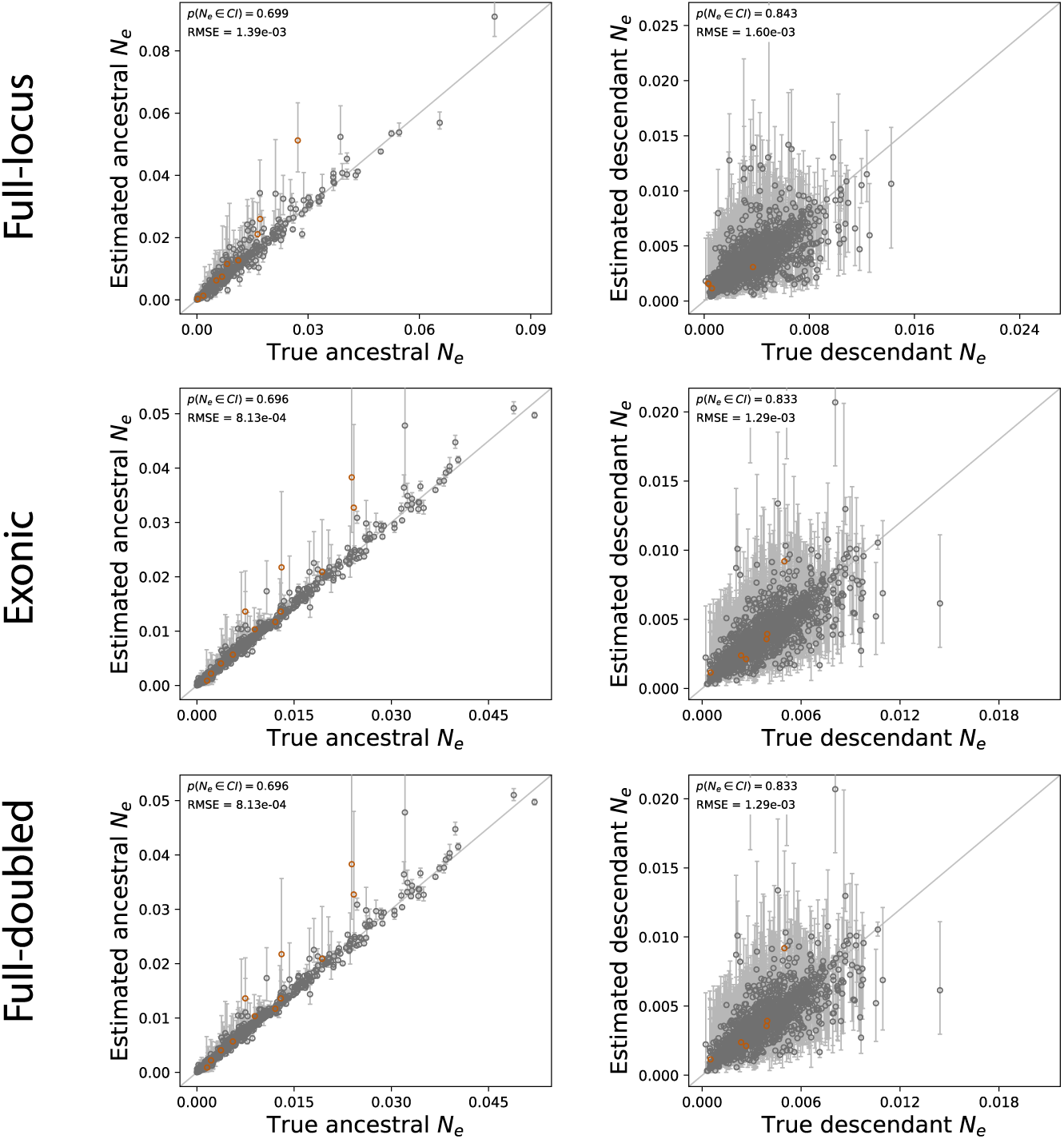
The accuracy and precision of effective population size estimates for ancestral populations (left column) and descendent populations (right column). The full-locus dataset is on the top row, exonic in the middle, and full-doubled on the bottom row. Each plotted circle and error bars represent the posterior mean and the 95% credible interval. Estimates for which the potential scale reduction factor was greater than 1.2 (indicating poor convergence/mixing) are highlighted in orange. Each plot contains 1,800 estimates (300 simulated datasets each with six effective population size estimates). Each plot shows the root-mean-square error (RMSE) and the proportion of estimates for which the 95% credible interval contained the true value – (*p*(*t* ∈ CI). Plots were generated using ‘pycoevolity’ v.0.2.4.

